# Vecmatch: Balancing Covariates in Multiple Groups Using Vector Matching Algorithm and Generalized Propensity Scores

**DOI:** 10.64898/2026.11.06.764786

**Authors:** Mateusz F. Kolek, Thomas Vogl

## Abstract

Covariate imbalance is a major source of bias in observational studies. This issue arises when confounding variables (e.g., age, sex) are distributed unevenly across compared groups, potentially influencing estimates of between-group differences in other dependent variables. Matching techniques based on propensity scores are widely used to address covariate imbalance and improve the accuracy of causal inferences. These methods work by calculating propensity scores based on imbalanced covariates and then matching observations with the closest propensity scores across compared groups. However, existing matching methods primarily focus on comparisons between at most two groups, and there is currently no widely implemented approach for matching across more than two treatment groups in popular programming languages.

In this paper, we present an R implementation of the vector matching algorithm for matching across multiple treatment groups. The implementation includes the core algorithm itself along with several extensions aimed at improving usability and flexibility in applied research. These features include tools for visualizing covariate distributions, functions for estimating generalized propensity scores, and methods for defining rectangular regions of common support. Users can choose among different matching methods and adjust various parameters to improve covariate balance. To support this process, we provide an optimization workflow based on the random search algorithm that identifies parameter combinations yielding the best matching quality and highest percentage of matched samples. In addition, the package includes functions for assessing post-matching quality, offering a comprehensive and flexible framework for robust causal inference in complex observational settings.

## 1. Introduction

Observational studies are inherently prone to various forms of bias, which can significantly influence the outcomes of subsequent statistical analyses. This may result in over- or underestimation of the true effects being studied, thereby reducing the internal validity of the research (Anderson et al., 2009; Grimes and Schulz, 2002). Causal inference from observational data is not inherently impossible, but it requires conditions, some of which are guaranteed by design in a randomized experiment, whereas in an observational study all of them must be assumed (Rubin, 1974, 1980; Herńan and Robins, 2020):

- **exchangeability** - the potential outcomes are independent of the treatment assignment given the measured covariates, *Y* (*t*) *⊥⊥ T | X* for every treatment level *t*,
- **positivity** - Pr(*T* = *t | X* = *x*) *>* 0 for every treatment level *t* and every covariate value *x* that occurs in the population,
- **consistency** - the observed outcome under a given treatment equals the counterfactual outcome under that treatment,
- **stable unit treatment value assumption (SUTVA)** - one individual’s outcome is unaffected by another individual’s treatment, and each treatment level corresponds to a single, well-defined intervention.

Exchangeability is most commonly threatened by confounding, which arises when a variable causally affects both the treatment assignment and the outcome. In this situation, the observed association is a mixture of the true causal effect and the confounder’s influence: depending on the direction of these associations, the resulting estimate can understate, overstate, or even reverse the sign of the true effect (Herńan and Robins, 2020).

The most fundamental method for eliminating confounding is randomization: the random assignment of individuals or objects to experimental conditions (Rubin, 1978). Randomization makes the potential outcomes independent of the treatment assignment, so that treatment groups are comparable in expectation with respect to both measured and unmeasured baseline characteristics. This is what makes treatment assignment ignorable (Gelman, 2007) and what establishes marginal exchangeability; assigning each treatment level with a known, nonzero probability establishes positivity at the same time. Consistency and SUTVA, by contrast, are not conferred by the assignment mechanism: the former requires each treatment level to correspond to a single, well-defined intervention (Herńan and Taubman, 2008), which in a trial is supplied by the protocol rather than by the randomization, and the latter can fail in a randomized experiment as readily as in an observational one whenever one subject’s treatment affects another’s outcome (Rubin, 1980; Hudgens and Halloran, 2008). In observational studies treatment is not randomized: exposure is not assigned by the researcher but reflects inherent characteristics (e.g. sex), self-selected behaviors (e.g. smoking), or clinical decisions, all of which may potentially be associated with the outcome. Randomization is not, however, the only route to exchangeability. Where treatment cannot be randomized, conditional exchangeability given a sufficient adjustment set can play the same role, provided the covariates responsible for confounding have been measured (Herńan and Robins, 2020). The question is therefore not whether causal effects can be estimated from observational data, but under what conditions, and by which methods, the required adjustment can be achieved.

Despite the complexities involved, a substantial methodological literature has developed over the past decades to address the problem of causal inference in observational settings (Anderson et al., 2009). One of the most intuitive approaches is to model the outcome conditional on the treatment and the measured confounders in a multivariate regression analysis (McNamee, 2005). Such outcome regression requires a correctly specified form of the relationship between the outcome and the confounders and sufficient overlap in the confounder space between the treatment groups. Even when both hold, the computed estimand is a conditional effect. It equals the corresponding marginal effect under an identity or log link with no treatment-by-covariate interaction, but not universally: for the logistic or Cox regression widely applied to biological data the adjusted coefficient is not equal to the marginal effect even in the absence of confounding, and its value depends on which covariates were included in the final model (Greenland et al., 1999; Martinussen and Vansteelandt, 2013). Obtaining a marginal effect from an outcome model requires predicting each subject’s outcome under every treatment level and averaging these predictions over the observed covariate distribution, which is also known as the g-formula, or g-computation (Snowden et al., 2011): an approach that remains substantially less used in practice than direct coefficient reporting (Vansteelandt and Keiding, 2011). The major issues of the outcome regression are that misspecification of the conditional mean translates directly into bias. Additionally the fitted model may extrapolate into regions of the covariate space in which one or more treatment groups are unrepresented. Both problems are difficult to detect from the fitted model alone (King and Zeng, 2006).

A second family of methods focuses on the covariate distribution. Subclassification and post-stratification analyze the treatment-outcome relationship within strata defined by the confounders and then pool the within-stratum estimates (Cochran, 1968). The approach, however, becomes impractical as the number of covariates grows, and is most natural for discrete confounders. Inverse probability weighting (IPW) achieves the same adjustment by modeling the probability of exposure rather than partitioning on it, weighting each unit by the inverse of its probability of receiving the treatment it actually received. It produces a pseudo-population in which treatment is independent of the measured covariates (Herńan and Robins, 2020). Because estimated probabilities close to zero or one yield extreme weights and unstable estimates, stabilized weights (Robins et al., 2000), truncation, and alternatives such as overlap weights (Li et al., 2018) are commonly applied. Extensions to multi-category treatments can be achieved by calculating generalized propensity scores described by Imbens (2000). Implementations for a wide range of estimators are available in the WeightIt package (Greifer, 2026).

A third group combines the methods used in the previous two, yielding estimators that remain consistent if either the treatment model or the outcome model is correctly specified. Augmented IPW (Bang and Robins, 2005) and targeted maximum likelihood estimation (Van Der Laan and Rubin, 2006) achieve this double robustness, and double/debiased machine learning (Chernozhukov et al., 2018) builds on the same estimating equations while allowing both nuisance functions to be estimated by flexible machine-learning methods.

A fourth approach mimics the randomization process. Instead of modeling the outcome or reweighting the sample, matching selects a subsample in which the covariate distributions of the treatment groups are similar (Cochran and Rubin, 1973; Rubin, 1973). It enforces common support, discarding units for which no comparable counterpart exists, thus eliminating the possibility of extrapolating over them. The cost is reduced sample size and an estimand defined on the matched, not the original sample. Matching on many covariates at once is made manageable by the propensity scores - the probabilities of treatment assignments given the measured covariates. They summarize the treatment-confounder relationship in a single scalar (Rosenbaum and Rubin, 1983, 1984).

Here and throughout, *treatment groups* denote any groups the researcher wishes to compare with respect to an outcome. They do not need to correspond to actual interventions. When the groups are defined by a condition rather than an assignable treatment, as in the example of Section 3, the contrasts defined in Section 2.1 compare the groups at a common covariate distribution. We then interpret them as covariate-adjusted differences rather than as effects of the condition, whose causal interpretation would additionally require well-defined versions of each level (Herńan and Taubman, 2008). Matching is also used for such non-causal comparisons, for example in studies of disparities (Stuart, 2010).

Developed originally for the binary design with one treatment and one control group, propensity scores were later generalized to multiple treatment values as the generalized propensity score (GPS) (Joffe and Rosenbaum, 1999; Imbens, 2000).

Matching becomes significantly more challenging in scenarios where multiple groups are present. In a basic binary groups study, matching involves finding pairs of observations with the closest propensity score values - this means the difference between propensity scores of the treatment and control should be minimized. However, when analyzing *k* groups, matching must be conducted based on a *k*-dimensional vector of GPS. This requires that the entire vector of *k* GPS values be similar between two observations for them to be considered a match. The challenge in this context arises from the absence of a single, easily computable measure to quantify the distance between these vectors. Unlike the binary case, where matching can be performed using the simple difference between propensity scores for the treatment and control groups, the multidimensional nature of *k*-group propensity scores complicates the assessment of similarity (see Figure 1).

**Figure 1:**
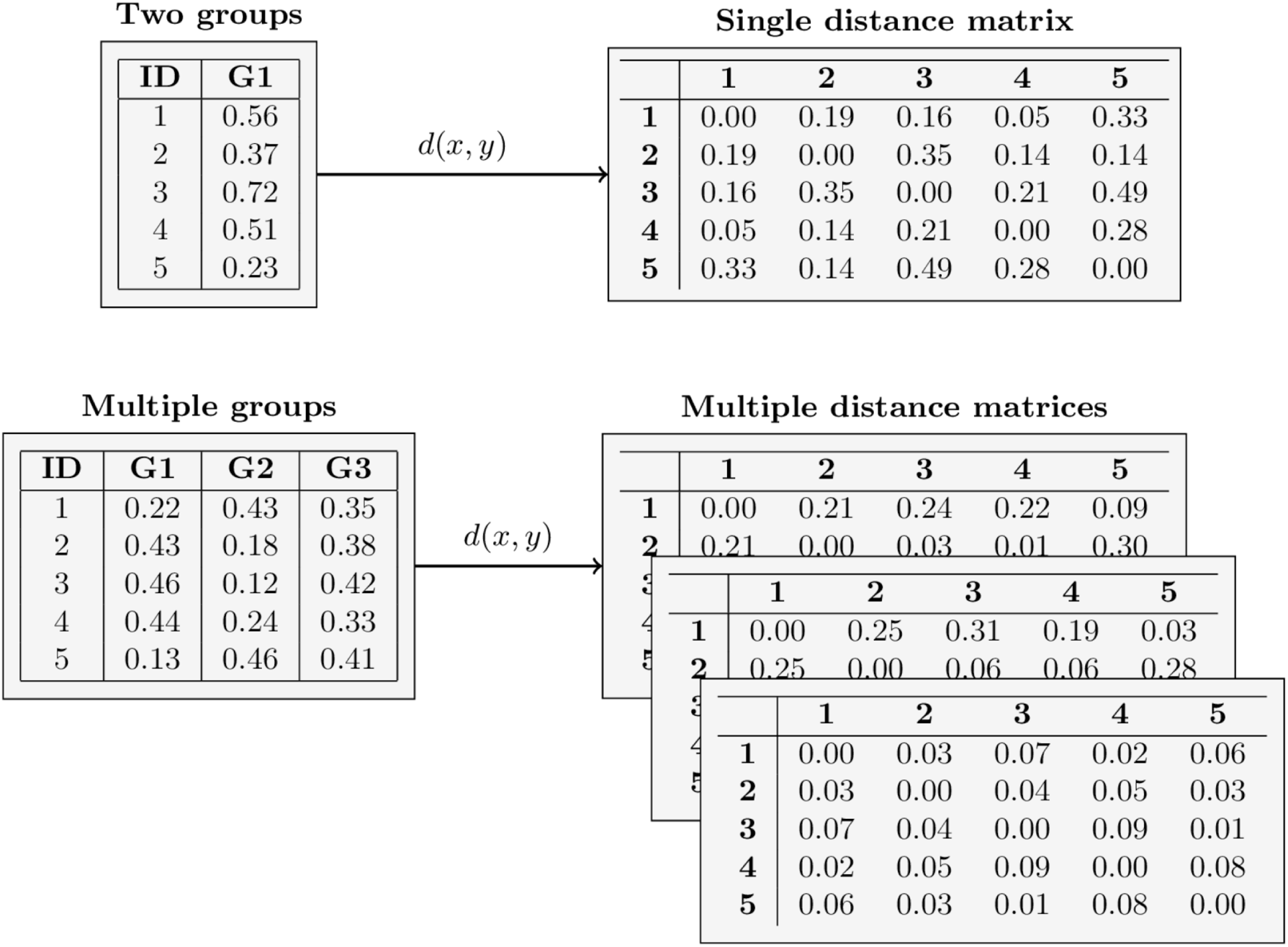
Illustration of the challenges involved in matching subjects across multiple treatment groups based on propensity scores. In the binary treatment case (top row), each subject is characterized by a single propensity score, representing the probability of assignment to one of two groups (with the second score being implicitly defined as 1 minus the first, due to the constraint that probabilities sum to 1). In this one-dimensional setting, Euclidean distances between scores can be directly computed to assess the similarity of subjects for matching. However, in the multi-group scenario (bottom row), each subject has a vector of propensity scores - one for each treatment group - resulting in multiple distance matrices. Separate distance matrices are derived for each score component, but there is no obvious scalar metric that captures overall similarity. As a result, a pair of subjects may appear to be close in one dimension but not in others, complicating the evaluation of match quality. The Euclidean distance is denoted in the diagram as *d*(*x, y*).

**Figure 2:**
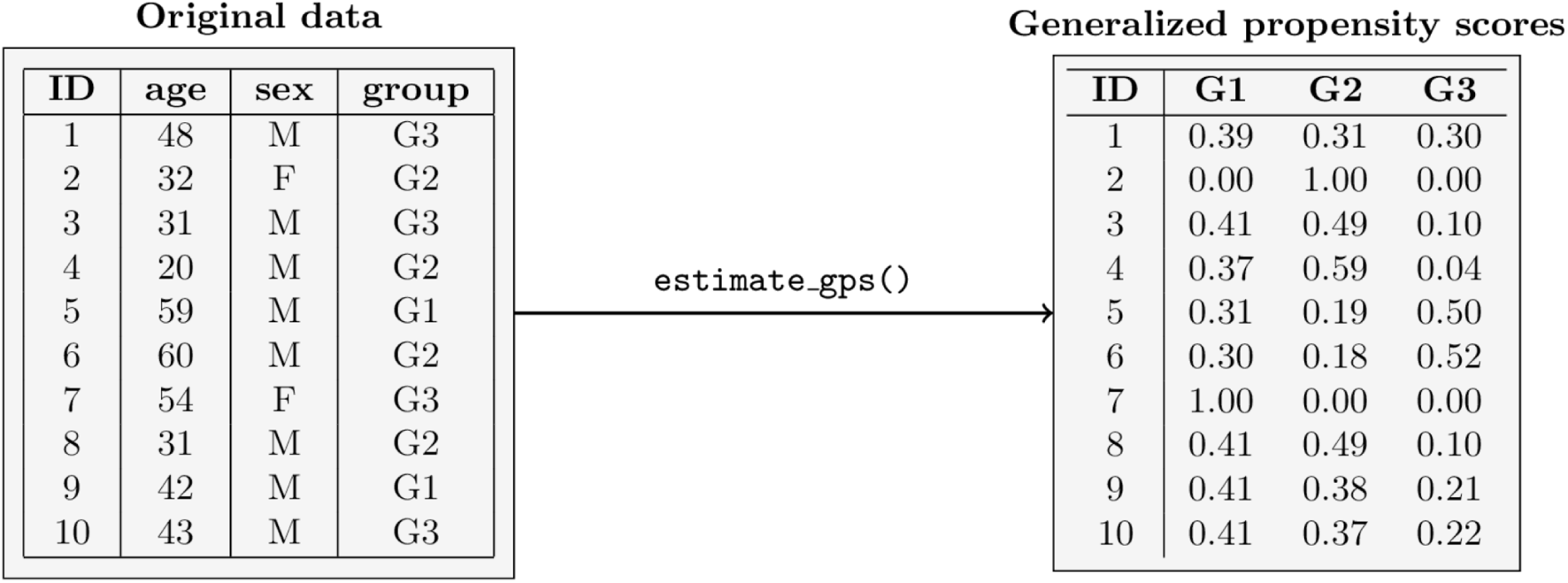
A visual representation of the generalized propensity score calculation process. For users new to the package, the `estimate_gps()` function can be understood as a black box that takes the original dataset - including covariates and treatment group assignments - as input and outputs a GPS matrix. A more detailed explanation of this transformation is provided in the accompanying text below.

An exemplary solution to this problem is presented by Lopez and Gutman (2017), where matching is combined with clustering of the GPS vector. The presented approach clusters observations with similar GPS values using the *k*-means method, and then performs matching within each cluster. It allows for matching only among observations whose entire GPS vectors are similar. The algorithm, referred to as *vector matching*, has not been available as a documented, maintained software package. Thus, the goal of the vecmatch package was to implement the method described in the article by Lopez and Gutman (2017), offering an easy-to-use workflow for matching observations across multiple treatment groups.

The development of vecmatch was motivated by a medical application in high-throughput Phage Immunoprecipitation Sequencing (PhIP-Seq). The high dimensionality in this setting is defined by the number of outcome variables, as a single assay measures antibody reactivity against hundreds of thousands of linear peptide antigens. The treatment variable is however low-dimensional and categorical. It typically represents a small number of clinically defined groups (e.g. healthy controls vs. three disease groups). The confounders are a handful of standard demographic and technical covariates such as age and sex, which are known to influence the outcomes (Vogl et al., 2021). The package therefore addresses the problem of a single multi-category treatment with a modest covariate set, not the substantially harder problem of high-dimensional treatments.

Two features make a matching-based approach particularly appropriate for PhIP-Seq applications. First, the outcomes do not exist at the point of design. Samples are selected from a pool of available biological material donors and then measured, so the design must be finalized and the samples must be selected before any outcome is available. The doubly-robust and machine-learning estimators discussed earlier cannot be used at the design stage to decide which samples to measure, since they require an outcome regression. The same constraint limits the practical value of weighting: weights require that every candidate sample has already been sequenced, whereas matching identifies the subset of samples to be measured in advance. In a cost-limited experiment the number of samples which can be measured is a major constraint, and can be easily controlled using matching. Second, the matched dataset is subsequently analyzed with established bioinformatics tools for enrichment calling, differential expression analysis and epitope-level differences. Several widely used tools provide no mechanism for sample weights or covariate adjustment, so confounding control must be achieved before the data enter the pipeline rather than within it. Constructing a balanced sample at the design stage delivers exactly this.

We therefore required a flexible and user-friendly workflow to match multiple cohorts based on several covariates, while still retaining direct access to the matched dataset for downstream bioinformatic analyses. The vecmatch package was developed to address this need.

## 2. Matching of multiple treatment groups

In this section, we will explore the statistical methods and approaches underlying the functions of the vecmatch package. We begin by defining the estimands that vector matching targets in the Target Estimand section (2.1). We then discuss the functions for generating graphical summaries and plots in the Data Visualizations section (2.2). Next, we will focus on the calculation of GPS in the Generalized Propensity Scores section (2.3) and Common Support Region section (2.4), and their matching process in the Vector Matching Algorithm section (2.5). We then examine various methods for evaluating the quality of matching in the Matching Quality Assessment section (2.6), and close with the Typical Workflow for Applied Users section (2.7).

### 2.1 Target estimand

Let *T ∈ T* = *{t*_1_*, …, t_k_}* denote the treatment variable, *X* the measured covariates, and *Y_i_*(*t*) the potential outcome of subject *i* under level *t*. Following Lopez and Gutman (2017), we assume that the estimands of interest are pairwise contrasts, this means comparisons of two single treatment levels rather than of groups of levels. In the simple binary case the two standard estimands are the average treatment effect (ATE) *E*[*Y_i_*(*t*_1_) *− Y_i_*(*t*_2_)] and the average treatment effect in the treated (ATT), *E*[*Y_i_*(*t*_1_) *− Y_i_*(*t*_2_) *| T_i_* = *t*_1_]. With *k* levels of the treatment variable the same definitions apply to each of the 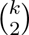 pairs, so the target is no longer a single number but a set of contrasts.

Vector matching enables estimating the *k −* 1 pairwise ATT-type contrasts relative to a reference level *t_c_*, restricted to subjects eligible for every treatment. Let *E_i_* = 1 indicate that subject *i* has all *k* GPS components inside the CSR borders defined in the Common Support Region section (2.4). The estimands are then

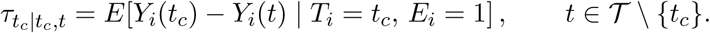

The role of vector matching is to make these contrasts estimable from the observed data. Under a regular assignment mechanism, subjects with the same GPS vector

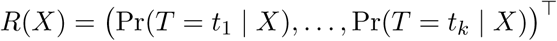

but different observed treatments are comparable, so differences in their outcomes can estimate the corresponding causal contrast (Imbens, 2000; Lopez and Gutman, 2017). It is worth noting that the equality of a single component Pr(*T* = *t | X*) is not sufficient: two subjects can share one component of the *R*(*X*) vector and differ in all others, and in such situation their outcome difference has no causal interpretation (Imbens, 2000). This explains why the vector matching algorithm clusters on the remaining GPS components before matching on the pair (*t_c_, t*), and why balance must be assessed across all arms simultaneously rather than pair by pair.

The requirement of joint eligibility has a second, practical consequence. If each pair were matched separately, as in a series of binary comparisons, each contrast would generalize to a different subset of the reference group, and the resulting effects would not be transitive: *τ_tc|tc,t_*_1_ and *τ_tc|tc,t_*_2_ could not be differenced to compare *t*_1_ with *t*_2_ (Lopez and Gutman, 2017). Conditioning every contrast on the same event *E_i_* = 1, and retaining in Step 5 only reference subjects matched in all arms, forces all the contrasts onto a single covariate distribution. They are then internally consistent and can be compared with one another. For assignable treatments, this is the multi-arm randomized trial that vector matching is intended to emulate. For groups defined by a condition, the matched contrast estimates the covariate-standardized difference

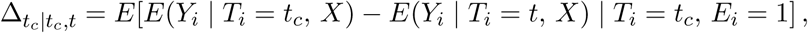

which compares each group with the reference group at the covariate distribution of the latter, and equals *τ_tc|tc,t_* only when the identification conditions of Section 1 hold.

### 2.2 Data visualizations

To enable users of the vecmatch package to quickly and comprehensively analyze the initial imbalance of the data, two types of plots were used: raincloud plots for continuous variables and mosaic plots for categorical variables. The raw plots can optionally be enhanced with summary statistics that numerically represent the extent of the imbalance.

#### 2.2.1 Raincloud plots

A raincloud plot (Figure 4) is essentially a composite plot consisting of several simple subplots that visualize the data distribution and key descriptive statistics derived from it (Allen et al., 2021). It includes:

- **half-split violin plots** - they illustrate the full probability density function (PDF) of the analyzed variable. They allow for easy identification of different distribution types (e.g., skewed, bimodal) across groups, enabling a preliminary assessment of imbalance
- **jittered point plots** - they display raw data as individual jittered points. Combined with violin plots, they help identify areas with the highest concentration of observations and detect potential outliers
- **boxplots** - they visualize key robust descriptive statistics, such as the median and quartiles
- **pointrange plots** - they represent the arithmetic means of individual groups using points, while standard deviations are shown with lines. They are overlaid on the violin plots

**Figure 3:**
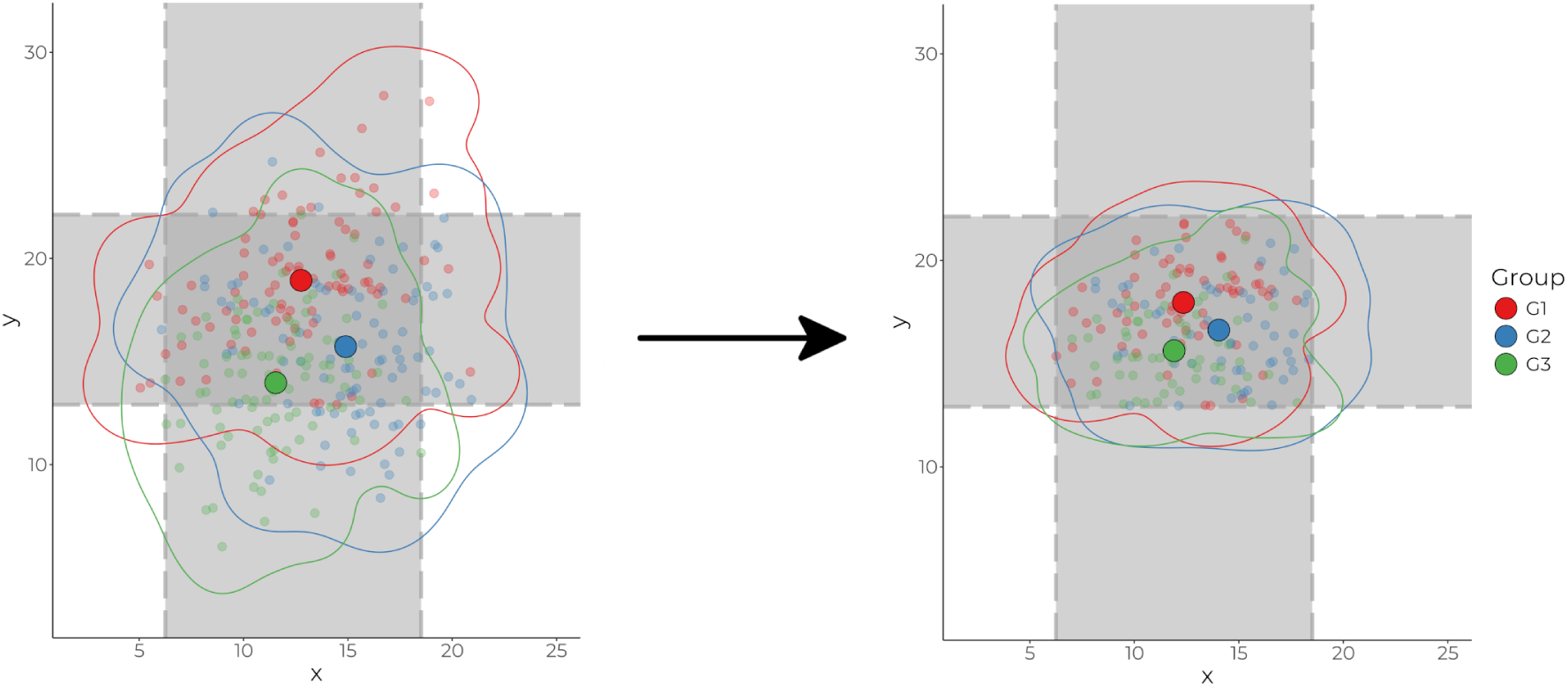
The distribution of data for two variables (x and y) across three exemplary groups (G1, G2, G3) is visualized using a scatterplot with color-coded groups. Individual observations are represented by colored points, while the 95th percentiles of the kernel density are illustrated with contour lines. Large points with black outlines indicate the arithmetic means of each group. Gray lines and shaded areas depict the CSR boundaries for each variable separately, whereas the rectangular region in the center of the plot represents the common support region (CSR) itself. On the left side of the plot, raw observations from a sample dataset generated using a bivariate normal distribution with different means and variance-covariance matrices for each group are displayed. On the right side, only the observations within the CSR are shown - all other observations have been removed.

**Figure 4:**
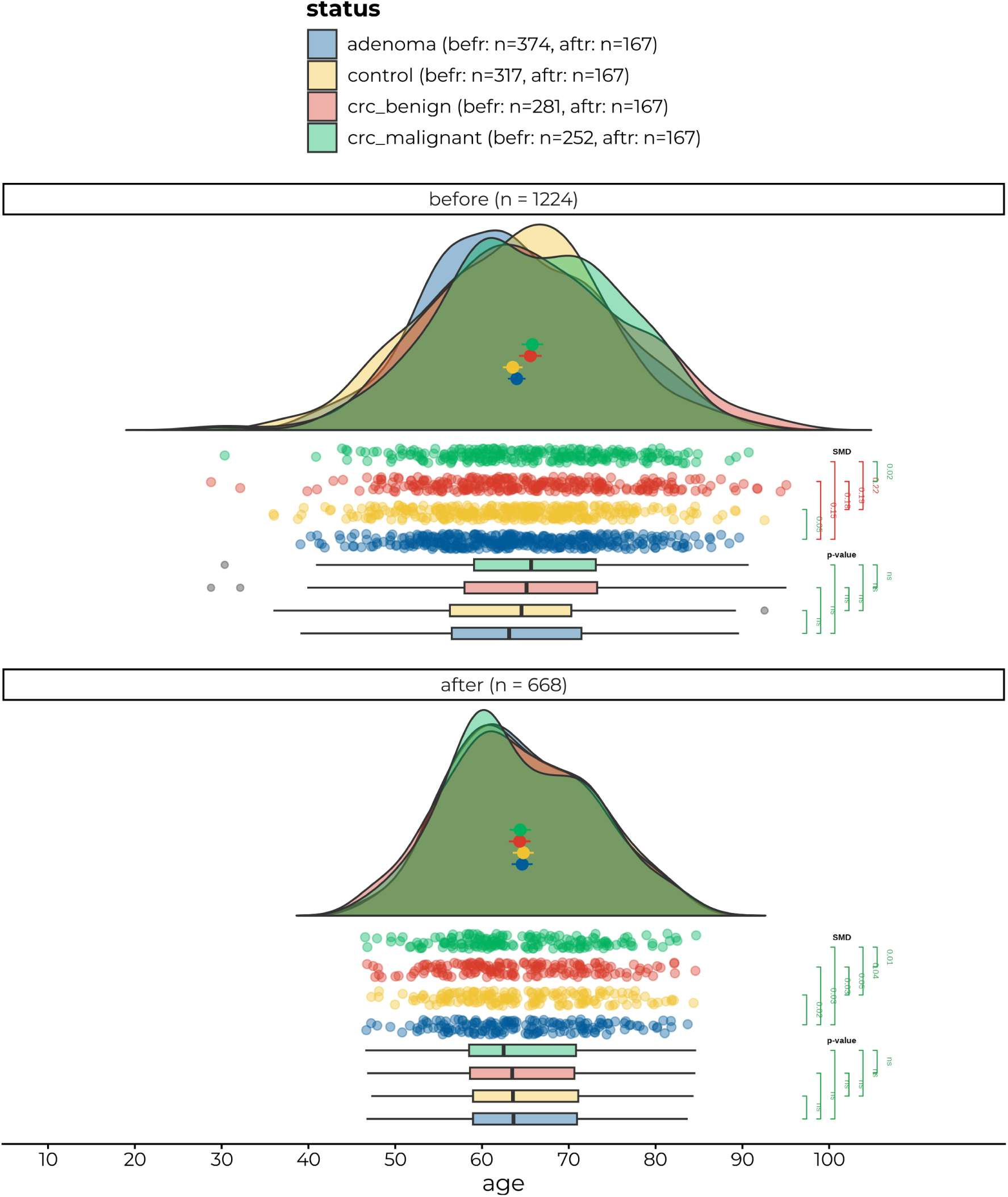
Conditional distribution of age across different status groups. The standardized mean differences (SMD) are represented by the brackets on the right side of the jittered point plot. The results of the Student’s *t*-tests are displayed alongside the boxplots on the right, offering insights into the statistical significance of the differences in age between the groups. The *n* in the facet labels indicates the total number of observations within each facet group, that is the full sample before matching and the matched sample after it. In the legend, the *n* values shown next to each status group label correspond to the counts before and after matching, respectively, following the alphabetical order of the facet labels.

To facilitate imbalance analysis, we added two types of statistics to the raincloud plot: the effect size coefficient and the statistical significance of the selected test. The type of test used in the analysis can be chosen using the significance argument, while the type of effect size can be specified with `smd_type`. Both parametric and non-parametric tests are available, all performed with a correction for multiple comparisons - the *p* values displayed to the user are already adjusted. For the `smd_type` argument, there are two options. The default value, `“mean”`, calculates Cohen’s *d* effect size coefficient. The alternative, `“median”`, computes the *r* effect size, derived from the *Z* statistic of the Wilcoxon test. The statistical significance values between two selected groups are displayed next to the brackets connecting them on the boxplot, while the effect size values are shown next to the jittered point plots. An example of a raincloud plot is shown in Figure 4.

#### 2.2.2 Mosaic Plots

A mosaic plot (Figure 5) is a type of visualization used to represent the relationship between two discrete variables. Its fundamental concept is to depict group frequencies using rectangles, where the area of each rectangle is directly proportional to the count of a given group in the contingency table (Wickham and Hofmann, 2011). In the case of the `mosaic()` function, the width of a rectangle is proportional to the frequency of the category on the x-axis, while its height corresponds to the frequency of the category on the y-axis. To ensure easy interpretation of the rectangle areas, the user can explicitly display the group counts on the plot by setting the `group_counts` argument to `TRUE`.

**Figure 5:**
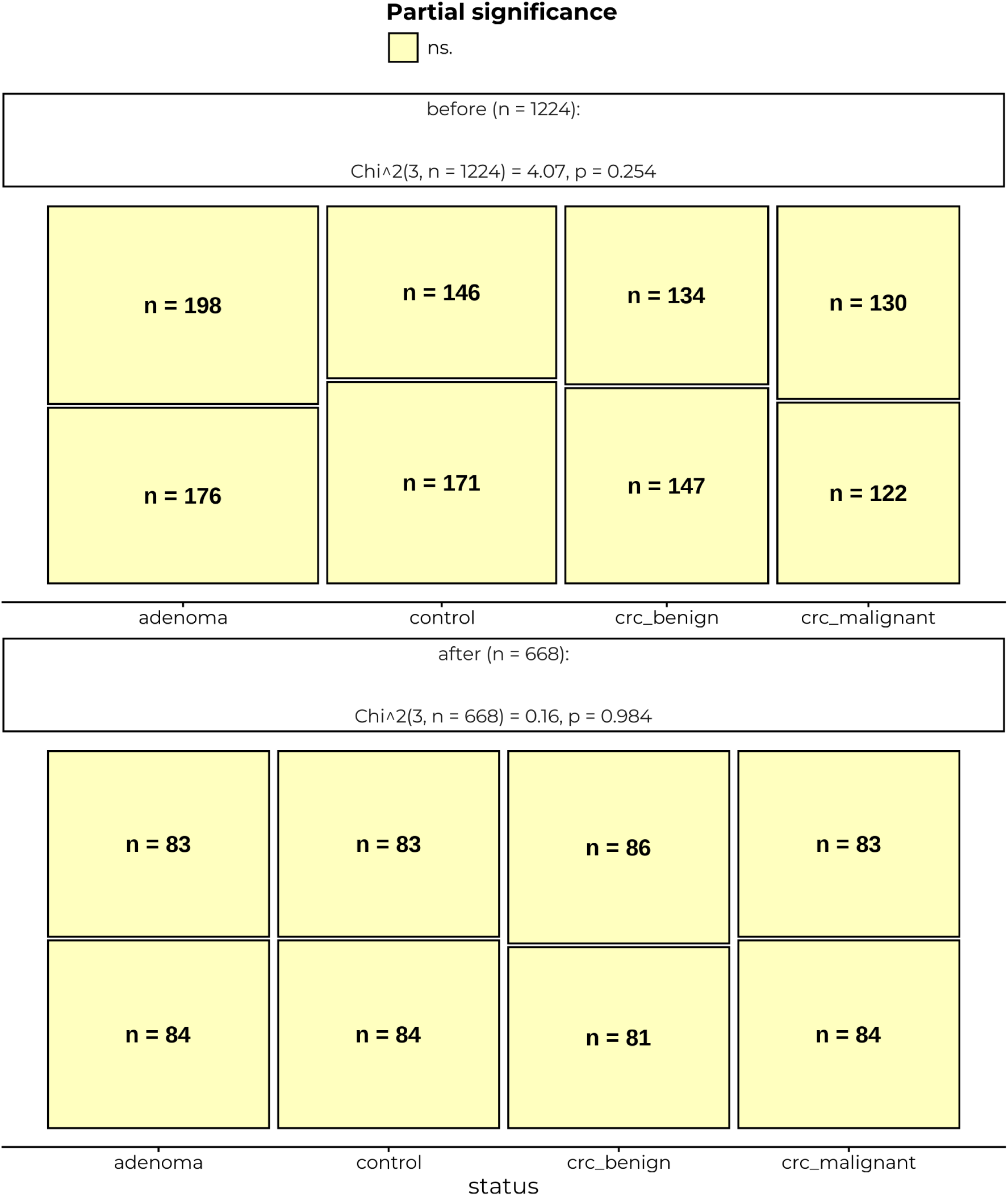
Conditional distribution of sex across different status groups. The counts for all groups are displayed inside the corresponding boxes, representing each subgroup within the mosaic plot. Partial significance is visually coded through color filling aesthetics. The labels provide the results of the Chi-squared test of independence, summarizing the statistical assessment of the relationship between the sex and status variables.

As noted by Hofmann (2003), there are several possible variations of mosaic plots depending on how the base contingency table is structured. To simplify the selection process for users, the `mosaic()` function adopts a fixed partitioning approach, where rectangles are always divided parallel to the y-axis. This means that for a given x-axis category, the rectangle widths remain the same across all y-axis categories, while the heights vary depending on the number of observations within each subgroup (see Figure 5, where, for example, the width of the `adenoma` category is the same for both women and men, while the heights differ based on the number of individuals in each status group). In our view, this approach significantly facilitates conditional comparisons of the distribution of the y-axis variable across different x-axis subgroups. For this reason, we recommend that users always place the treatment variable on the x-axis and the predictor variable, for which imbalance is being assessed, on the y-axis.

The `mosaic()` function also provides the option to perform a Chi-square test of independence and report its results in the plot’s footer when the `significance` argument is set to `TRUE`. Additionally, the rectangles representing group frequencies are filled with colors based on standardized Pearson residuals. Since standardized adjusted Pearson residuals are normally distributed with a mean of zero and standard deviation of 1 by definition, they can be used to determine the partial significance of the cell counts. The color cutoffs were chosen based on the most commonly used significance levels *α*: 0.050 for residuals between 1.96 and 2.58, 0.010 for residuals between 2.58 and 3.29, 0.001 for residuals greater than 3.29. As the distribution of the residuals is symmetrical, for negative Pearson residuals, the corresponding negative cutoffs were applied.

### 2.3 Generalized propensity scores

Propensity scores can be defined as the treatment assignment probabilities, representing the likelihood that an observation belongs to a specific treatment group given the observed confounders (Gelman, 2007). In other words, they quantify how likely it is that a given subject (e.g., patient or sample) is assigned to a particular group based on the observed covariates included in the model. Propensity scores are a fundamental component of the matching process, as matching is typically performed using these scores or GPS in the case of multiple treatment groups. GPS summarize the measured covariates included in the model in a single vector of values, allowing for comparison across observations. At a given value of the score vector, the distribution of the measured covariates is the same in every treatment group (Rosenbaum and Rubin, 1983; Imbens, 2000), even though individual subjects with similar scores may differ in their covariates.

Formally, if we define the treatment group of interest as *t*, the set of all possible treatments as *T*, where *t ∈ T*, and the vector of covariates as *X*, the propensity score can be expressed as Pr(*t | X*) (Imbens, 2000). Generalized propensity scores extend this concept by computing assignment probabilities for each treatment level separately, resulting in a vector of treatment assignment probabilities (compare Figure 1) rather than a single value, as in scenarios with binary treatment (Lopez and Gutman, 2017; Imbens, 2000).

For example, consider Table 1, which presents ten observations, each assigned to one of three treatment groups, as indicated by the group variable. The generalized propensity scores are computed using the original dataset on the left that includes all covariates along with the treatment (group) assignment. In this case, the model formula used is group ∼ age * sex, meaning that both age, sex, and their interaction are used to predict group membership. The resulting GPS matrix has dimensions 10 *×* 3, reflecting the ten observations and the three unique treatment groups. For each observation (i.e., each row in the matrix), a vector of three probabilities is calculated, representing the estimated likelihood of assignment to each treatment group given the covariates. The probabilities in each row sum to 1 because the treatment groups are mutually exclusive and exhaustive. The ultimate purpose of the vector matching algorithm is to identify observations whose entire GPS vectors (i.e., entire rows in the matrix) are most similar to one another, ensuring comparability across groups based on the joint distribution of the covariates.

**Table 1:** Generalized propensity scores example data.

| ID | age | sex | group |
|---|---|---|---|
| 1 | 48 | M | G3 |
| 2 | 32 | F | G2 |
| 3 | 31 | M | G3 |
| 4 | 20 | M | G2 |
| 5 | 59 | M | G1 |
| 6 | 60 | M | G2 |
| 7 | 54 | F | G3 |
| 8 | 31 | M | G2 |
| 9 | 42 | M | G1 |
| 10 | 43 | M | G3 |

**Table 2:** Sample of the cancer dataset with 10 observations. The cancer dataset is a synthetically generated dataset, whose design was based on metadata used for matching patients for subsequent PhIP-Seq analyses. It contains 5 variables and 1224 observations. The variable representing treatment status is labeled status and includes 4 unique categories: control, representing healthy controls; adenoma, for patients diagnosed with colorectal adenomas during colonoscopy; crc_benign, for patients with early-stage (localized) colorectal cancer; and crc_malignant, for patients with advanced colorectal cancer (the level names crc_benign and crc_malignant are retained from the package data). The remaining variables (sex, age, bmi, and smoker) differ across the status groups, which motivates matching. In practical applications, after matching, PhIP-Seq analyses would be performed to compare the antibody profiles of patients across the groups. Because disease status is a condition rather than an assignable treatment, the matched comparison in this example is descriptive: it compares antibody profiles across disease groups at a common age and sex distribution.

| status | sex | age | bmi | smoker |
|---|---|---|---|---|
| adenoma | M | 74.79 | 28.08 | no |
| adenoma | F | 57.05 | 22.00 | no |
| crc_malignant | M | 68.47 | 26.13 | no |
| crc_malignant | M | 75.04 | 26.27 | no |
| adenoma | F | 74.13 | 22.25 | no |
| adenoma | M | 56.18 | 26.18 | yes |
| crc_malignant | F | 58.48 | 26.02 | no |
| adenoma | M | 70.01 | 31.52 | yes |
| crc_benign | M | 73.09 | 24.18 | no |
| crc_malignant | M | 83.76 | 23.99 | yes |

Various models can be used to estimate the GPS in a multiple treatment groups scenario. For nominal treatments, where the order of the treatment groups does not matter, multinomial models can be applied, such as multinomial logistic regression models (Venables and Ripley, 2002), vector generalized linear models for multinomial data (Yee, 2015), multinomial response models with bias reduction using the Poisson trick (Kosmidis et al., 2020), or baseline category logit models (Elff, 2022). For ordinal treatment variables, a proportional odds logistic regression model can be used (Venables and Ripley, 2002). The vecmatch package integrates these models from the mentioned packages, enabling users to estimate GPS with a single function: estimate_gps. Users can also utilize other functions, as well as machine learning and deep learning algorithms that are not currently supported by the package. However, in such cases, the GPS matrix must be created manually.

### 2.4 Common support region

Matching observations is meaningful only when their GPS values lie within a common probabilistic space - in other words, they have a common range of GPS. This space in matching theory is defined as the common support region (CSR - see Figure 3) and it is the overlap area of the distributions of the GPS in all treatment groups (Caliendo and Kopeinig, 2008). Following the approach of Lopez and Gutman (2017), the vecmatch package implements the calculation of the CSR based on the maximum and minimum GPS values within individual treatment groups. The boundaries of the CSR determined in this way take a rectangular shape (as shown in Figure 3). All observations that fall outside the defined CSR boundaries are removed from further analysis. Lopez and Gutman (2017) recommend re-estimating the GPS after excluding observations outside the CSR boundaries. To account for changes in the sample after applying CSR filtering, vecmatch allows users to re-estimate the GPS by setting the argument `refit = TRUE in the csregion()` function. This triggers a re-fit of the original GPS model using the exact same specification as in the previous `estimate_gps()` call, but restricted to the subset of data within the defined CSR.

### 2.5 Vector matching algorithm

The vector matching algorithm, designed by Lopez and Gutman (2017), enables matching based on the GPS across multiple treatment groups. It utilizes the *k*-means method to cluster observations with similar GPS values and then performs matching within these clusters, pairing observations with comparable GPS values. Matching is conducted in pairs, using the control group and each of the treatment groups included in the study. At the final stage, after completing the matching process, a merge of all matched observations is performed. The resulting matched dataset includes only those control group observations that have been matched with at least one observation from each treatment group, along with their respective matches.

A detailed description of the vector matching algorithm can be found in Lopez and Gutman (2017). Below, we aim to provide a more accessible explanation of the vector matching algorithm for non-technical users by translating the mathematical notation used in the original work. Steps 1, 2, and 3-5 of the algorithm are implemented using the functions estimate_gps, csregion, and match_gps, respectively. Users should execute these functions sequentially, one after another in the indicated order, to perform the complete matching procedure. The package follows a modular design, allowing users to independently adjust key parameters at each stage of the process, thereby offering flexibility and fine-grained control over the matching procedure.

We briefly outline the vector matching algorithm implemented in the vecmatch package and illustrate each step using the cancer dataset, which contains four treatment groups: healthy controls, colorectal adenomas, and early-stage and advanced colorectal cancers.

1. **Estimate the vector of generalized propensity scores (GPS).** For each subject, estimate the GPS vector using a multinomial regression model or another appropriate statistical model. In vecmatch, this is performed by the `estimate_gps()` function, which returns a matrix with one column per treatment group. In the cancer dataset, the resulting GPS matrix has four columns, corresponding to the four treatment groups, and 1224 rows, corresponding to unique patients.
2. **Restrict to the common support region (CSR).** Next, determine the boundaries of the CSR and remove all subjects whose GPS values fall outside this region. In vecmatch, this step is performed by the `csregion()` function. Optionally, the GPS model from Step 1 can be refitted on the restricted sample using the same formula and model specification. For the cancer dataset, the CSR calculation leads to the exclusion of some observations whose GPS vectors are incompatible with at least one of the treatment groups, thereby improving the overlap between groups.
3. **Apply a logit transformation to the GPS values.** To improve numerical stability and the behavior of distances in subsequent steps, the GPS values are transformed using the logit function: 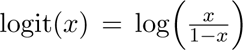, where *x* denotes a single treatment assignment probability in the GPS matrix. In vecmatch, this transformation is performed automatically inside `match_gps()`. In the cancer example, the entire four-column GPS matrix is thus transformed before clustering and matching are carried out.
4. **Select a control group and perform clustering-based matching.** If no natural control group is present, choose as reference the group whose average GPS value lies closest to the center of the distribution across all treatment groups. Denote the chosen control group by *t_c_*and any other treatment group by *t_i_*. For each *t_i_*not equal to *t_c_*, proceed as follows:

1. *Cluster on the remaining GPS components.* Remove from the transformed GPS matrix the columns corresponding to *t_c_*and *t_i_*. Perform *k*-means clustering on the remaining columns to obtain *K* clusters of observations (the default in vecmatch is *K* = 5). This step groups subjects with similar GPS vectors for the remaining treatment groups. In the cancer dataset, we set *t_c_* = control and apply *k*-means clustering separately for each non-control group (adenoma, crc_benign, crc_malignant).
2. *Match within clusters on the control and current treatment.* After clustering, discard the GPS columns used for clustering and focus only on the logit-transformed GPS components for *t_c_* and *t_i_*. Within each of the *K* clusters, perform 1:1 matching between subjects in the control group and subjects in the current treatment group. Matching can be carried out with or without replacement, using algorithms such as *k*-nearest neighbors or full optimal matching, and an optional caliper can be specified. In the cancer example, subjects assigned to a given cluster (e.g., cluster 1) are only allowed to match with other subjects from the same cluster. This restriction encourages similarity of the entire GPS vector, rather than just the two components corresponding to *t_c_* and *t_i_*. The procedure is repeated iteratively for each non-control group (adenoma, early-stage and advanced colorectal cancer).
5. **Construct the final matched dataset.** After repeating Step 4 for all non-control treatment groups, collect all control subjects that have been matched at least once, along with their matched counterparts. When multiple treatment groups are present, one can optionally restrict the final dataset to those control observations that appear in all pairwise matched sets. In the cancer case study, this yields three intermediate matched datasets: controls vs. adenoma, controls vs. early-stage colorectal cancer, and controls vs. advanced colorectal cancer. The final analysis dataset is then constructed by retaining only those control subjects that are present in all three matched datasets, together with their corresponding matches in each treatment group.

### 2.6 Matching quality assessment

#### 2.6.1 Standardized mean differences

The standardized mean difference (SMD), also referred to as standardized bias (Lopez and Gutman, 2017; Rubin, 2001), represents a simplified adaptation of Cohen’s *d* effect size coefficient. It is commonly used in propensity score matching to evaluate the balance of covariates across treatment groups. The SMD is computed separately for each covariate and for every possible pairwise combination of treatment groups, as defined by the following equation:

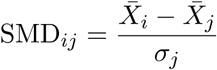

where SMD*_ij_* denotes the standardized mean difference between groups *i* and *j*, *X̅_i_* and *X̅_j_* represent the mean values of covariate *X* in groups *i* and *j*, respectively, and *σ_j_* is the standard deviation of *X* within group *j*. Each unordered pair is evaluated once, in the order in which the treatment levels appear in the factor, so the denominator is the standard deviation of the second level of the pair rather than a pooled standard deviation.

Following the computation of the SMD for all pairwise group comparisons, the absolute values are taken, and the maximum SMD is selected, in accordance with the methodology proposed by Hade (2012) and Lopez and Gutman (2017). Additionally, the average SMD can be calculated for each covariate, an approach implemented using the parameter statistic = “mean” in the vecmatch package.

The literature presents a range of recommended thresholds for determining sufficient matching quality, with values ranging from 0.25 (Rubin, 2001) to 0.1 (Nguyen et al., 2017). However, Monte Carlo simulations conducted by Nguyen et al. (2017) indicate that adopting a more strict threshold of 0.1 is preferable to minimize bias in treatment effect estimation.

#### 2.6.2 Percentage of matched samples

This metric provides a straightforward and intuitive measure of matching quality, indicating the proportion of observations from the original dataset that were successfully included in the matched dataset. While no specific threshold exists for this metric, the proportion of matched observations is highly dependent on the initial imbalance of the dataset.

It is worth noticing that there is a significant difference between the implementation in the vecmatch package and the one proposed by Lopez and Gutman (2017). In vecmatch, it represents the percentage of the original observations that appear in the matched dataset. For instance, if the original dataset contains 1,200 observations after applying the `csregion()` function, and the percentage of matched samples is 72 percent, then the final matched dataset consists of 1,200 * 72 percent = 864 observations. In contrast, Lopez and Gutman (2017) defines it as the percentage of control observations from the original dataset that are retained in the matched dataset. Since vector matching with the *k*-nearest neighbors algorithm (with replacement) prioritizes maximizing the number of controls, the values in Lopez and Gutman (2017) were usually consistently high across all simulations and examples (above 80 percent). However, this is not necessarily the case in our implementation, as the percentage of matched observations in other treatment groups is typically lower than in the control group.

#### 2.6.3 Characterizing the matched population

Both diagnostics above describe how similar the treatment arms are to one another before and after matching. They say nothing about how similar the matched sample is to the sample that the user started with, and this distinction matters for interpretation. As discussed in Section 15.4 of Herńan and Robins (2020), matching on propensity scores produces an effect for the subgroup of units picked out by the score values used in matching. That subgroup is a product of the GPS model, the CSR borders, the caliper and the matching method, and it is not chosen by the analyst in advance. A matched sample can be perfectly balanced and still represent a narrow and unfamiliar slice of the original population or even the original sample used for matching.

The population is defined by the distribution of the propensity scores, but it does not need to be described by them. Because the GPS is a function of the measured covariates, the retained subjects can be characterized directly in the covariate space, which is the scale on which the reader can judge whether the matched sample still represents the population of interest. We therefore encourage users to report the covariate distributions before and after matching alongside the balance statistics. Three functions in vecmatch support this. `raincloud()` and `mosaic()` both accept a facet variable, so binding the original and the matched data into a single data frame and faceting on it (see the Step 5: post-matching quality assessment section (3.5)) places the two distributions side by side for continuous and categorical covariates, respectively. `balqual()` reports the per-group counts before and after matching, from which group-specific retention can be read directly. It also accepts the mutually exclusive type = “desc_full” and type = “desc_reduced”, either of which can accompany the standard balance metrics in the same function call. Both argument values describe the covariates as they appear in the formula. They follow the layout of the balance tables, with the statistics in the rows and the matching stages in the Before and After columns. For numeric covariates, “desc_full” reports the number of observations, mean, standard deviation, minimum, quartiles, median, maximum, and the moment estimators of skewness and excess kurtosis, while “desc_reduced” keeps only the number of observations, minimum, mean, median and maximum. For categorical covariates, each level is cross- tabulated as a count with its percentage computed for each variable, separately before and after matching. Comparing the Before and After columns within a treatment level shows how far, and in which direction, the retained subpopulation has moved, whereas comparing treatment levels at a fixed stage shows the balance itself. Substantial shifts in the mean or quantiles, a marked compression of the range, or a change in the level percentages indicate that the matched population differs from the original.

### 2.7 Typical workflow for applied users

In practice, an applied user typically proceeds as follows. First, generalized propensity scores are estimated using `estimate_gps()`, which summarizes covariate information into one or more scores guiding the matching procedure. Second, the common support region is determined with `csregion()`, which may remove a non-negligible fraction of observations but ensures that only comparable subjects are retained. Third, matching is performed via `match_gps()`, yielding a matched dataset whose quality can be assessed using standardized mean differences and the proportion of matched samples. Finally, more elaborate workflows, including optimization of tuning parameters, are illustrated in the package vignettes accompanying vecmatch.

## 3. Practical example

In this section we will describe an exemplary data analysis on the cancer dataset from the vecmatch package, matching multiple cohorts based on selected covariates. The vecmatch package has a strict workflow, following the vector matching algorithm defined by Lopez and Gutman (2017) and it is advisable to follow it. The whole process consists of 5 steps and ensures the best possible matching quality using the vector matching algorithm.

### 3.1 Step 1: data exploration and initial imbalance assessment

The vecmatch package provides two graphical functions for assessing initial dataset imbalance: `raincloud()` and `mosaic()`. The `raincloud()` function is designed for continuous variables, while the `mosaic()` function is intended for usage with categorical variables. Both functions allow grouping by up to two categorical variables (using the group and facet aesthetics) and calculate various statistical summaries, including significance tests and effect size coefficients, as discussed in previous sections.

In this exemplary analysis, we focus on two predictor variables from the cancer dataset: the discrete sex and the continuous age. To evaluate differences in age across status groups, we can use the `raincloud()` function as follows:

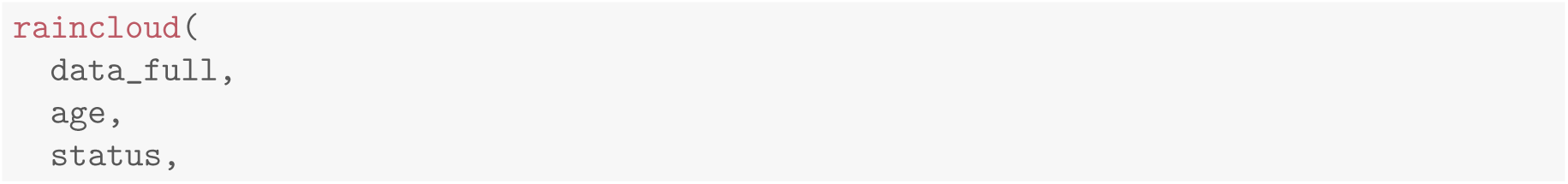
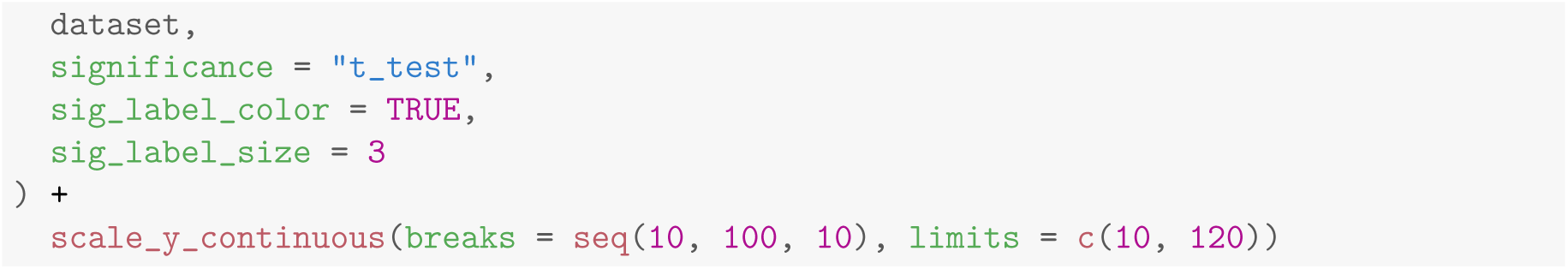

This code will generate the plot shown in Figure 4.

Similarly, we can evaluate the differences in the sex variable using the `mosaic()` function. The code snippet below demonstrates this:

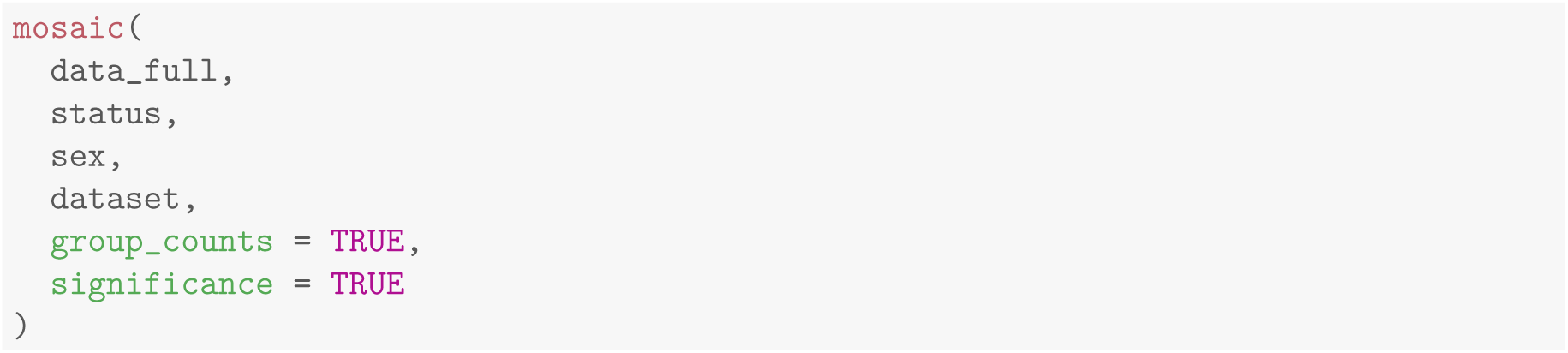

Note that both plots are generated using the data_full data frame, which stacks the full original sample (before) and the matched sample (after) - see the Step 5: post-matching quality assessment section for reference. The values discussed in this step are those of the before facet.

Based on the plots, we can observe imbalance in the age (Figure 4) and sex (Figure 5) variables. For age, four of the six standardized mean differences (SMDs) exceed the 0.10 threshold, the largest reaching 0.22, so the status groups are clearly imbalanced. The Student’s *t*-tests reported alongside the plot are adjusted for multiple comparisons and none of them is significant, but this does not establish balance. The power of such a test depends on the sample size, whereas balance is a property of the realized sample rather than a hypothesis about a population, so balance should be judged on the SMDs and the tests read as an exploratory summary only (Imai et al., 2008).

After a closer examination of the mean age values, we can notice that they form two separate clusters. The first cluster includes controls and patients with adenomas, who tend to be younger, while the second cluster consists of patients with early-stage and advanced colorectal cancer, who are generally older. The primary objective of the matching process will be then to align these two clusters, ensuring they have a common mean age value.

For sex, three of the six SMDs exceed the 0.10 threshold, the largest reaching 0.14. The overall Chi-square test of independence and the partial tests displayed in Figure 5 are not significant, but, as for age, that reflects the size of the sample rather than the balance of the groups.

Both covariates therefore carry imbalance of the kind the vector matching algorithm is intended to remove.

### 3.2 Step 2: estimation of generalized propensity scores

The next step in the vector matching algorithm is the calculation of generalized propensity scores (GPS) for the treatment variable. These scores represent the treatment assignment probabilities and are based on user-defined covariates. The relationship between the treatment variable and predictors can be easily defined using R-specific formula notation, allowing for both additive and interaction effects.

The vecmatch package provides a straightforward method for calculating the GPS matrix with different methods using the `estimate_gps()` function. However, custom approaches can also be applied within the vector-matching workflow. The GPS matrix used in subsequent analyses must have the exact same structure as the output of estimate gps, containing the treatment variable and the treatment assignment probabilities for each level of the treatment variable. Importantly, the probabilities for each row in the GPS matrix must sum to 1. The resulting data frame has to be of class gps.

In the following example, we will use a simple formula with age and sex as predictors, allowing for an interaction effect. The method used to estimate the generalized propensity scores (GPS) will be a multinomial logistic regression, which can handle multiple levels of the treatment variable. The code for estimating the GPS using the multinomial logistic regression model is as follows:

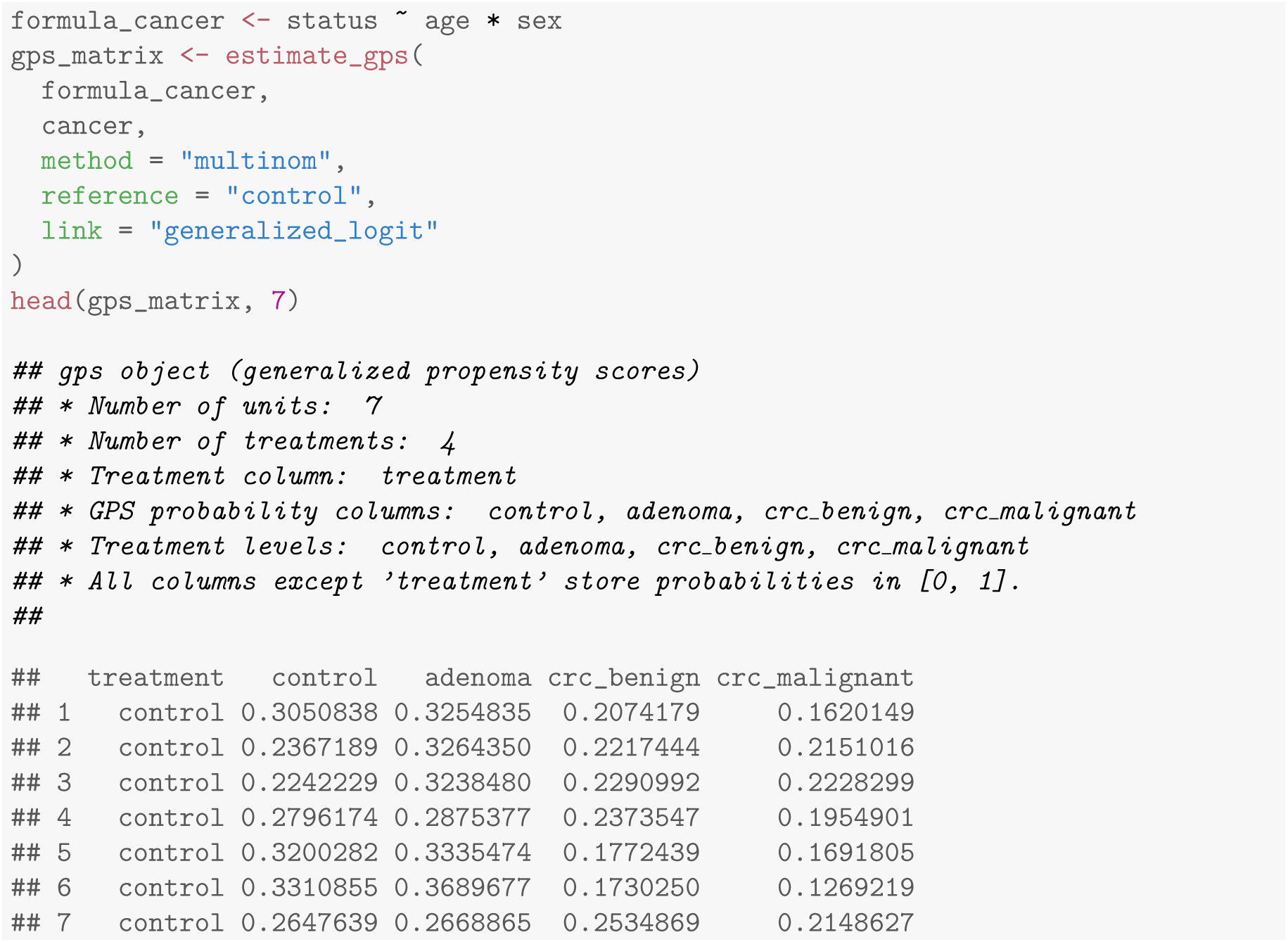

### 3.3 Step 3: calculating common support region borders

To remove observations that are not eligible for further matching, the borders of the common support region (CSR) must be calculated. The `csregion()` function facilitates this process by taking the gps_matrix object as its sole argument. It computes the CSR borders and automatically filters the input gps_matrix to include only the observations that fall within the calculated CSR borders.

The function returns an object of class csr, which contains the filtered data along with several additional attributes. These attributes enable users to manually filter the initial gps_matrix or access a summary of the CSR border calculation as a data frame object. The `csregion()` function can be used as follows:

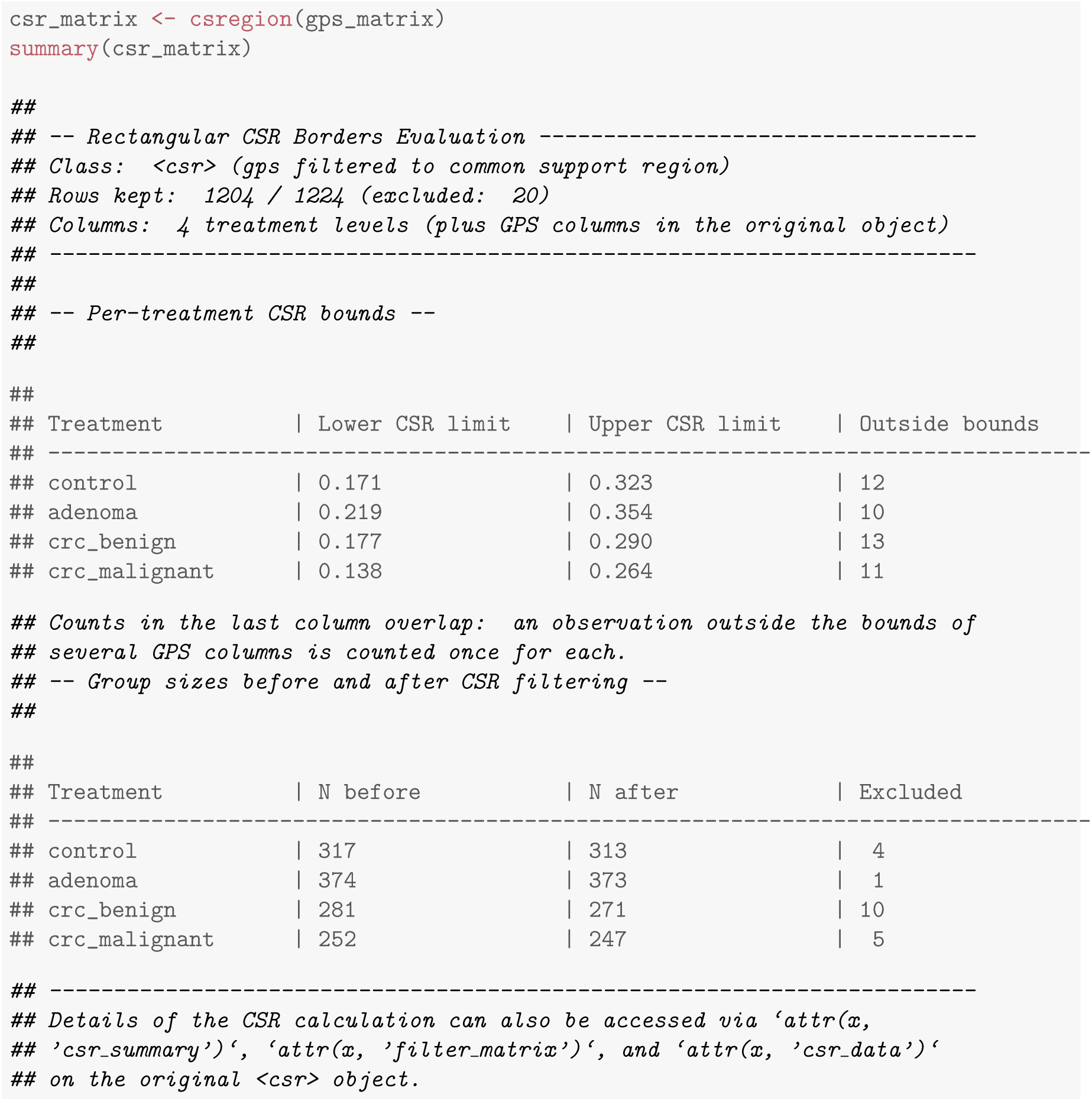

We can then compare the dimensions of the original gps_matrix and filtered csr_matrix.

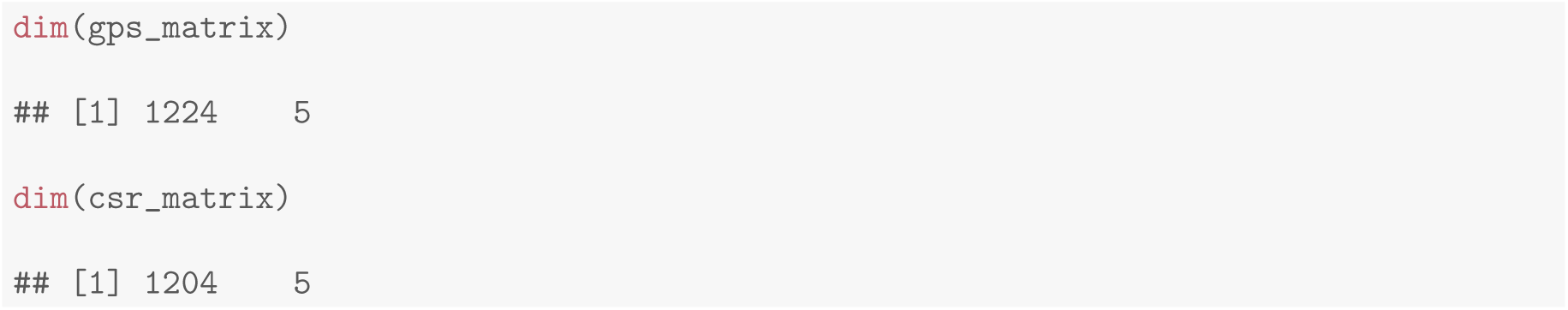

Clearly, 20 observations have been filtered out of the original matrix.

Next, as recommended by Lopez and Gutman (2017), the GPS is recalculated using only the data within the CSR. The `csregion()` function performs this refitting by default, reusing the formula and the estimation method of the original `estimate_gps()` call, so the csr_matrix object above already contains the refitted scores. Setting refit = FALSE skips the refitting and returns the original scores restricted to the CSR.

### 3.4 Step 4: *k*-means clustering and matching

The most crucial step of the vector matching algorithm is the *k*-means clustering and subsequent matching within the resulting clusters. This approach ensures that only observations with similar GPS vectors are matched, thereby justifying the name *vector matching*. The core components of this step are the `stats::kmeans()` function (R Core Team, 2024), which performs clustering, and the `Matching::Matchby()` or `optmatch::fullmatch()` functions (Sekhon, 2011; Hansen and Klopfer, 2006), which match observations within each *k*-means cluster. These functionalities are combined into a single function named `match_gps()`.

The `match_gps()` function takes the csr_matrix as input and returns a matched dataset. The matching process can be customized using various arguments, and it is advisable to experiment with different parameter combinations to optimize both matching quality and the number of matched samples. For the replicability of results, it is also crucial to set the random number generator seed before running the clustering and matching procedures. This ensures that the clustering assignments and matches are consistent across different runs of the analysis, allowing others to reproduce the results exactly.

The caliper argument deserves particular attention here, as it governs both the quality of the matching and the number of retained observations, and its effect is not monotone for most matching methods. Readers interested in how the caliper behaves across its range, and in how to select a value for their own data, are referred to the Appendix section on caliper width (C). The value used below was obtained by applying exactly the procedure described there: the matching was repeated over a grid of caliper values with all other arguments held fixed, the maximum SMD was plotted against the percentage of matched observations, and among the values meeting the strict balance criterion of 0.10 the one maximizing retention in a stable region of the curve was selected. The number of *k*-means clusters is left at the package default of five, so that the clustering constraint that defines vector matching is in force.

The match `gps()` function can be used as follows:

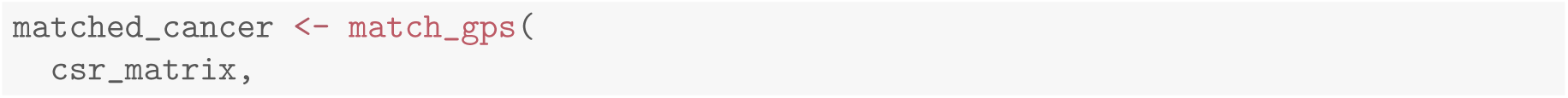
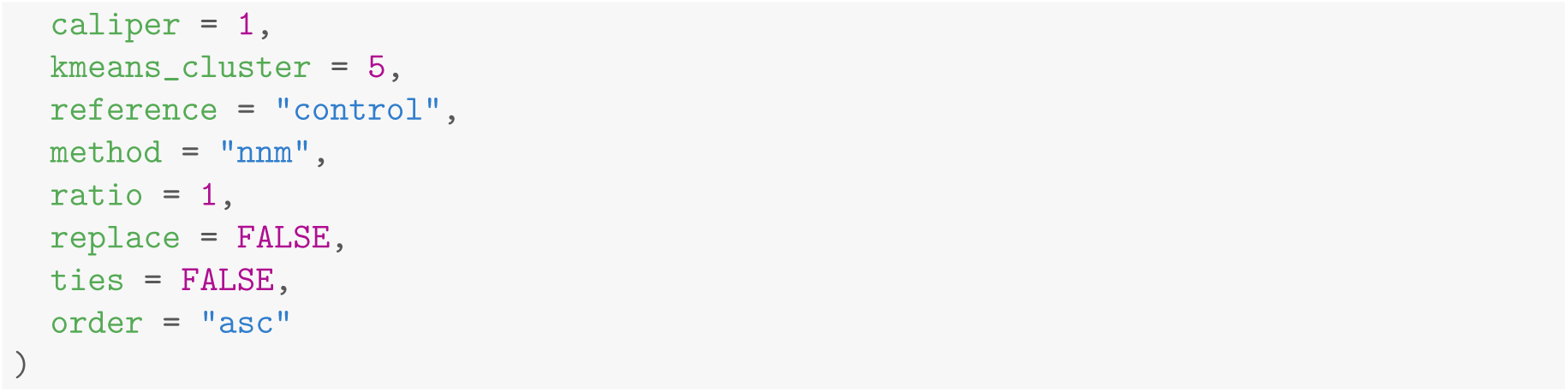

### 3.5 Step 5: post-matching quality assessment

The vecmatch package provides a dedicated function, `balqual()`, to assess the quality of post-matching results. This function compares various metrics and statistical summaries between the pre- and post-matching datasets, using a user-defined formula to evaluate balance. Its usage is straightforward and requires only the matched dataset (produced by `match_gps()`) and the formula used for the calculation of the GPS. Besides the balance metrics themselves, the type argument controls which summaries are returned: setting it to “desc_reduced”, as below, additionally prints the descriptive statistics introduced in Section 2.6.3, computed separately for the pre- and post-matching samples, so that the effect of matching on the covariate distributions can be inspected directly alongside the standardized mean differences:

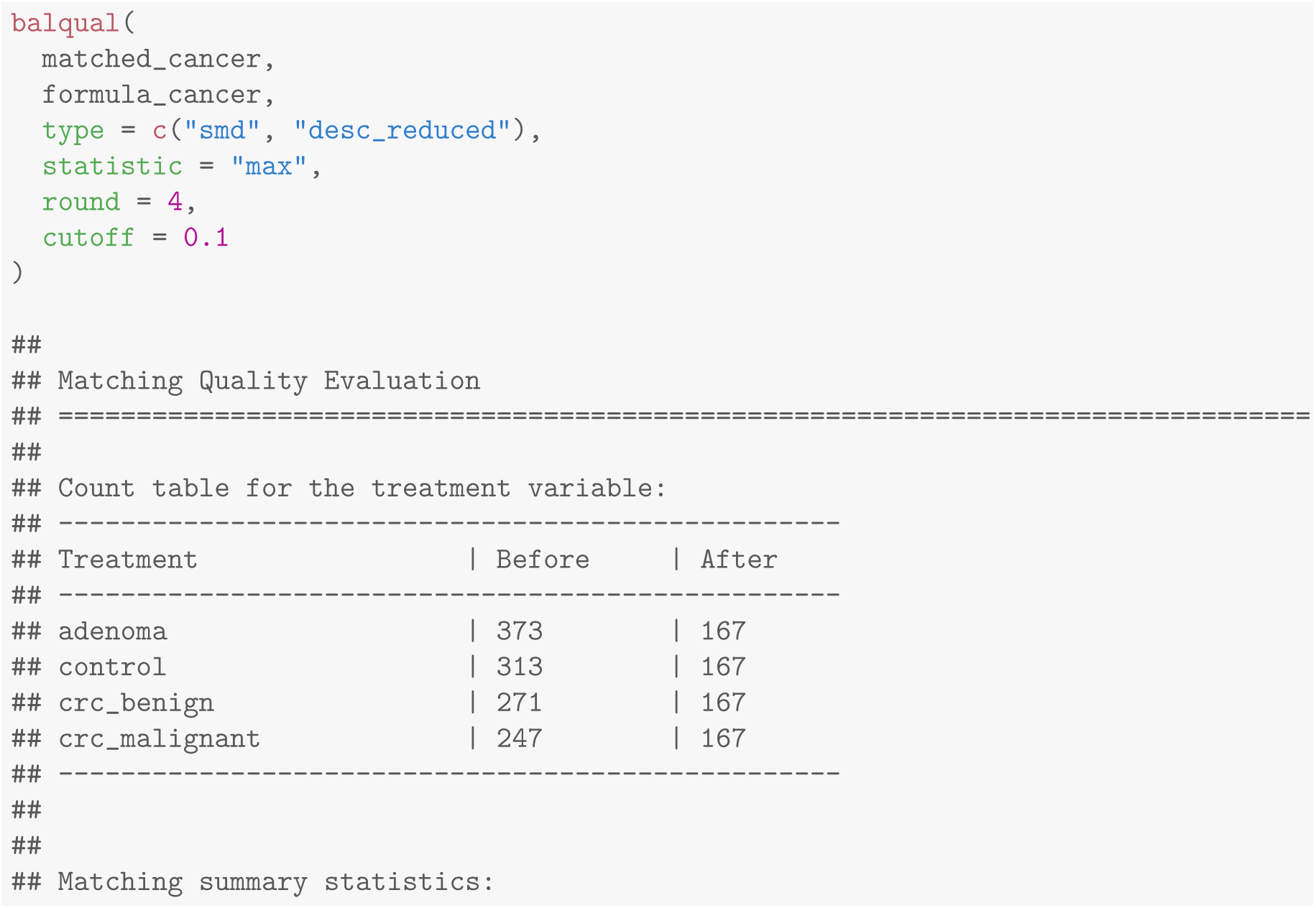
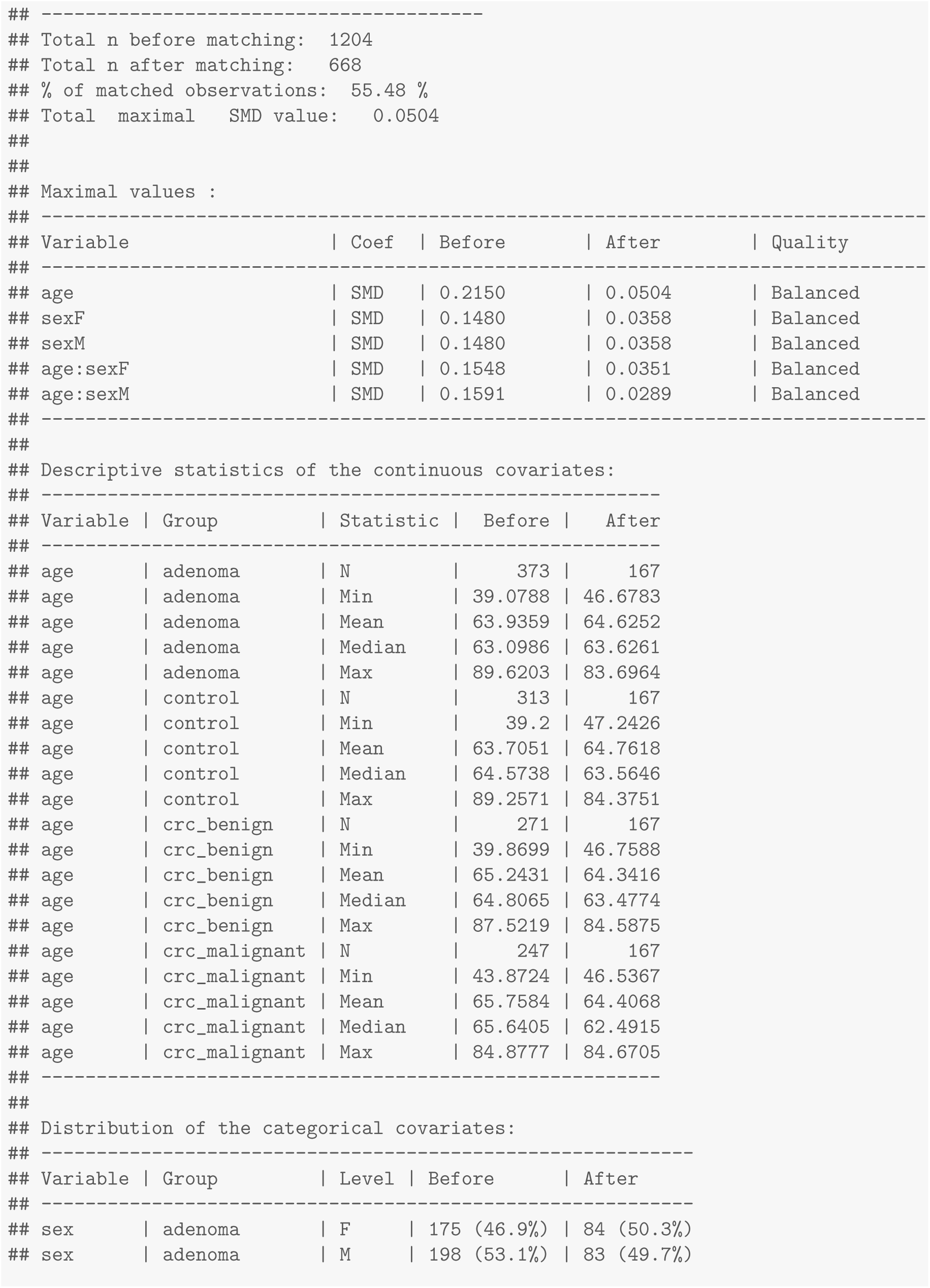
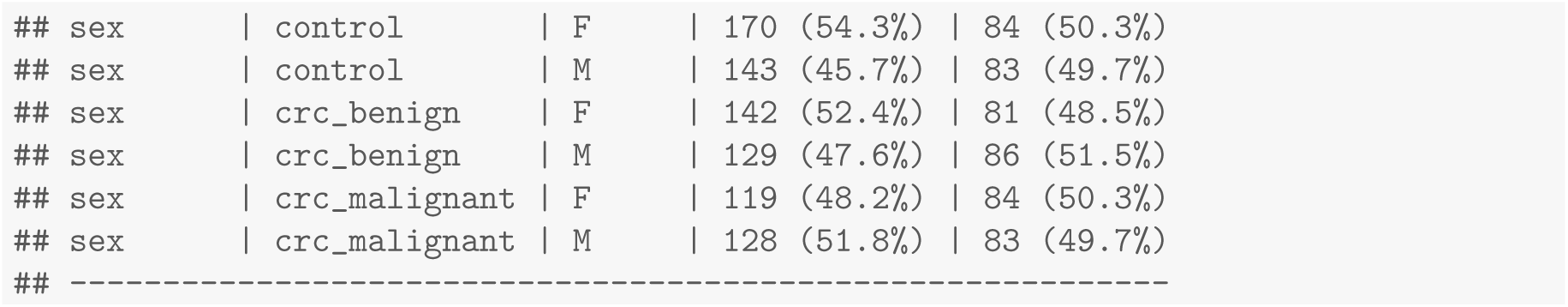

After assessing the quality with `balqual()`, we can stack the sample we started from and the matched sample into a single data frame to visualize age and sex before and after matching using `raincloud()` and `mosaic()`. Note that the before facet is the full original sample, not the observations discarded by the matching, so the two facets are nested rather than disjoint:

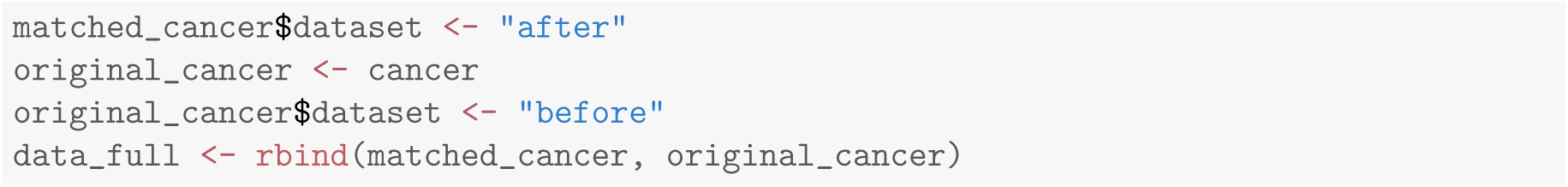

The corresponding raincloud and mosaic plots are shown in Figure 4 and Figure 5, respectively.

## 4. Summary and discussion

The vecmatch package provides a structured and flexible framework for matching across multiple treatment groups, implementing the vector matching algorithm proposed by Lopez and Gutman (2017). It is one of the few available R packages designed for multi-group matching. In contrast, widely used packages such as MatchIt (Ho et al., 2011), optmatch (Hansen and Klopfer, 2006), Matching (Sekhon, 2011), and twang (Cefalu et al., 2024) are primarily intended for binary comparisons between one treatment and one control group. Although TriMatch supports up to three groups (Bryer, 2017), it is no longer actively maintained. The closest existing alternative is multilevelMatching (Yang et al., 2016), which implements the matching and stratification estimators of Yang et al. (2016) for three or more treatment levels on a scalar function of the GPS. It targets the same design problem as vecmatch, but provides matching estimators of the pairwise effects rather than a modular workflow that returns the matched dataset itself, and it offers no tooling for common support, balance diagnostics, visualization or parameter tuning. A related but distinct effort is CausalGPS (Khoshnevis et al., 2023), which implements matching and weighting on the generalized propensity score for continuous exposures rather than for multiple discrete groups. It shares the use of the GPS as the basis for matching with vecmatch, but targets a dose-response relationship instead of pairwise contrasts between treatment arms. As a result, researchers working with multicategory treatments often resort to alternative methods such as weighting or subclassification. The vecmatch package aims to fill this methodological gap by offering a dedicated, up-to-date solution for multi-group matching.

vecmatch is built with modularity in mind. It allows users to separately handle the estimation of generalized propensity scores (GPS), define the common support region (CSR), and perform the matching itself using customizable methods. In addition to the default *k*-nearest neighbor matching with replacement, the package includes full optimal matching (Hansen and Klopfer, 2006), which often improves the percentage of matched observations. The modular design also enables fine-grained control over the matching parameters at each stage.

Despite its flexibility, the matching process in vecmatch - as with all propensity score- based methods - has important limitations. One key limitation is that matching necessarily excludes some observations to ensure comparability across groups. This reduction in sample size is influenced by several factors, including the initial covariate imbalance, the number of treatment groups, and the relative group sizes. Our experience further indicates that the choice of matching algorithm and its parameters can substantially affect the proportion of matched observations and overall balance.

In practice, researchers may prefer to retain all observations in certain treatment groups, particularly when dealing with rare diseases, costly or hard-to-obtain samples, or predefined cohorts. However, we stress that it is neither statistically valid nor scientifically defensible to fix the number of matched observations in a particular group. First, observations that fall outside the common support region must be excluded to satisfy the assumption of overlap - a core requirement of the Rubin-Neyman causal inference framework (Rubin, 1974). Violating this assumption by retaining unmatched samples compromises the validity of the matching and leads to extrapolation, rendering subsequent causal inferences unreliable.

Second, even when all observations fall within the CSR, the matching algorithms implemented in vecmatch do not support constraining the number of retained samples in specific groups. Enabling such constraints would require significant alterations to the underlying algorithms and would likely only be feasible in simpler two-group scenarios.

Nonetheless, to address practical concerns about sample retention, vecmatch includes an optimization workflow. This feature uses a random search algorithm to identify hyperparameter combinations that maximize the number of matched observations in selected groups while still satisfying the CSR condition. This approach provides a principled balance between preserving rare samples and maintaining statistical validity.

Another challenge users may face is the complexity of tuning numerous hyperparameters. These settings can strongly influence both the quality of the match (e.g., standardized mean differences) and the percentage of observations retained. To assist users in navigating this complexity, vecmatch includes dedicated functions that help optimize parameter choices in a data-driven way.

Finally, it is important to recognize that the limitations of vecmatch are not unique to the package but arise from foundational principles in observational causal inference. The requirement for a common support region implies that meaningful comparisons can only be made when treatment groups share similar covariate distributions. If substantial differences exist - such as between elderly and young individuals or between genders - then matching may not be feasible, and causal claims cannot be made without risking bias.

### 4.1 Limitations of propensity-score matching and their relevance to GPS

King and Nielsen (2019) argue that propensity scores should not be used for matching. Their argument has three parts. First, PSM targets the wrong experimental analogue: matching on a scalar score approximates a completely randomized experiment, in which covariates are balanced only in expectation, instead of a fully blocked one, in which they are balanced exactly. Second, as a consequence, PSM exhibits the PSM paradox: beyond a certain point, pruning further units increases imbalance and model dependence, because within the region where the score is already balanced, further pruning is effectively random with respect to the covariates. Third, since many specifications remain defensible on the standard balance diagnostics while yielding different matched samples, an analyst who can observe the outcome estimate while choosing the matching parameters may select among them on the basis of the outcome, and the published output gives the reader no way to detect this.

None of these arguments is specific to the binary case, and all carry over to the GPS setting. Two qualifications apply to the first: the *k*-means step restricts matches to subjects with similar values on the remaining GPS components, imposing a coarse blocking constraint absent from scalar PSM, and King and Nielsen (2019) themselves accept propensity scores for removing extreme common-support violations, which is the role of `csregion()` here.

The paradox is directly observable from the package’s own output. Read from high to low retention, Figure 8 is the diagnostic plot: an interval over which the maximum SMD rises as the percentage of matched observations falls is the paradox in this dataset. It is visible for full optimal matching and for KNNM with replacement, and absent for KNNM without replacement over the range examined. Users should therefore never adopt a single caliper (including the package default) without sweeping a grid and inspecting this plot, manually or through `optimize_gps()`. Two further checks follow. Because vector matching balances a *k*-dimensional score, balance must be assessed in the covariate space: `balqual()` reports SMDs for every term of the GPS formula and every pairwise contrast, and a match that balances the scores while leaving formula terms above the cutoff is an instance of the first concern. And because pruning redefines the population as well as the balance, the output of balqual(type = “desc_full”) and the faceted `raincloud()` and `mosaic()` plots should be read alongside the SMDs (Section 2.6.3 and 4.2).

The third argument has less power in the setting that motivated the package. As described in Section 1, matching in PhIP-Seq applications is a sample-selection step: subjects are drawn from a pool of available biological material and the assay is run only on the matched subset. At the point where `match_gps()` is called, no antibody reactivity has been measured and the outcome data do not exist. The specification is fixed before any effect estimate can be computed - not by the analyst’s restraint, but by the order in which the experiment is carried out - and any design that selects a matched sample in advance of measurement enjoys the same protection. The interface reinforces this: no function in the workflow accepts an outcome variable, and the optimization of Appendix B maximizes retention subject to a balance criterion, never an effect size.

This is a property of a design and an interface, not a guarantee about practice. A researcher applying vecmatch retrospectively has the discretion King and Nielsen (2019) describe, and we recommend two safeguards. The specification, including the caliper sweep of Appendix C and the balance criterion of Section 4.2, should be settled before the outcomes are examined. And the sweep should be reported rather than only its winner: a caliper presented alone cannot be distinguished from one selected post hoc, whereas the trade-off curve shows which specifications were considered and whether the reported one sits in a stable region.

### 4.2 Diagnosing failed matching

A matched dataset should not be judged on a single number. In our view, a well-performed match requires three conditions to hold at the same time, and vecmatch reports each of them separately.

1. **Sufficient retention.** The `csregion()` function reports how many observations fall inside the CSR borders, and `balqual()` reports the total counts before and after matching together with the percentage of matched observations. Because attrition is rarely uniform across arms, the per-group Before and After counts printed by `balqual()` are more informative than the overall percentage: a treatment group retained at a small fraction of its original size cannot support the contrasts in which it appears, however favorable the overall figure may look. We would treat a group retained at well under a fifth of its original size as a warning sign rather than apply any fixed threshold, since the tolerable loss depends on the initial group sizes and on the precision required.
2. **Acceptable balance.** The `balqual()` function with type = “smd” reports the mean and maximum standardized mean differences across all pairwise contrasts and all terms of the model formula, and flags the result against a user-defined cutoff. Balance must hold across all arms simultaneously, not pair by pair, for the reasons given in the Target Estimand section (2.1).
3. **A matched population that still resembles the original one.** The type = “desc_full” and type = “desc_reduced” arguments of `balqual()` describe the covariates before and after matching, and the facet argument of `raincloud()` and `mosaic()` displays the same comparison graphically. As discussed in the Characterizing the Matched Population section (2.6.3), a sample can be perfectly balanced and still describe a narrow and unfamiliar slice of the original data.

These three diagnostics are not interchangeable, and a match that fails any one of them is not a successful match, even when the other two appear satisfactory. We stress that all three are produced by the package itself, from a single `balqual()` call supplemented by the two plotting functions, so every failure mode discussed below can be identified from the package’s own output without additional tooling. Table 3 summarizes the combinations we consider most common in practice, together with the output in which each is visible.

**Table 3:**
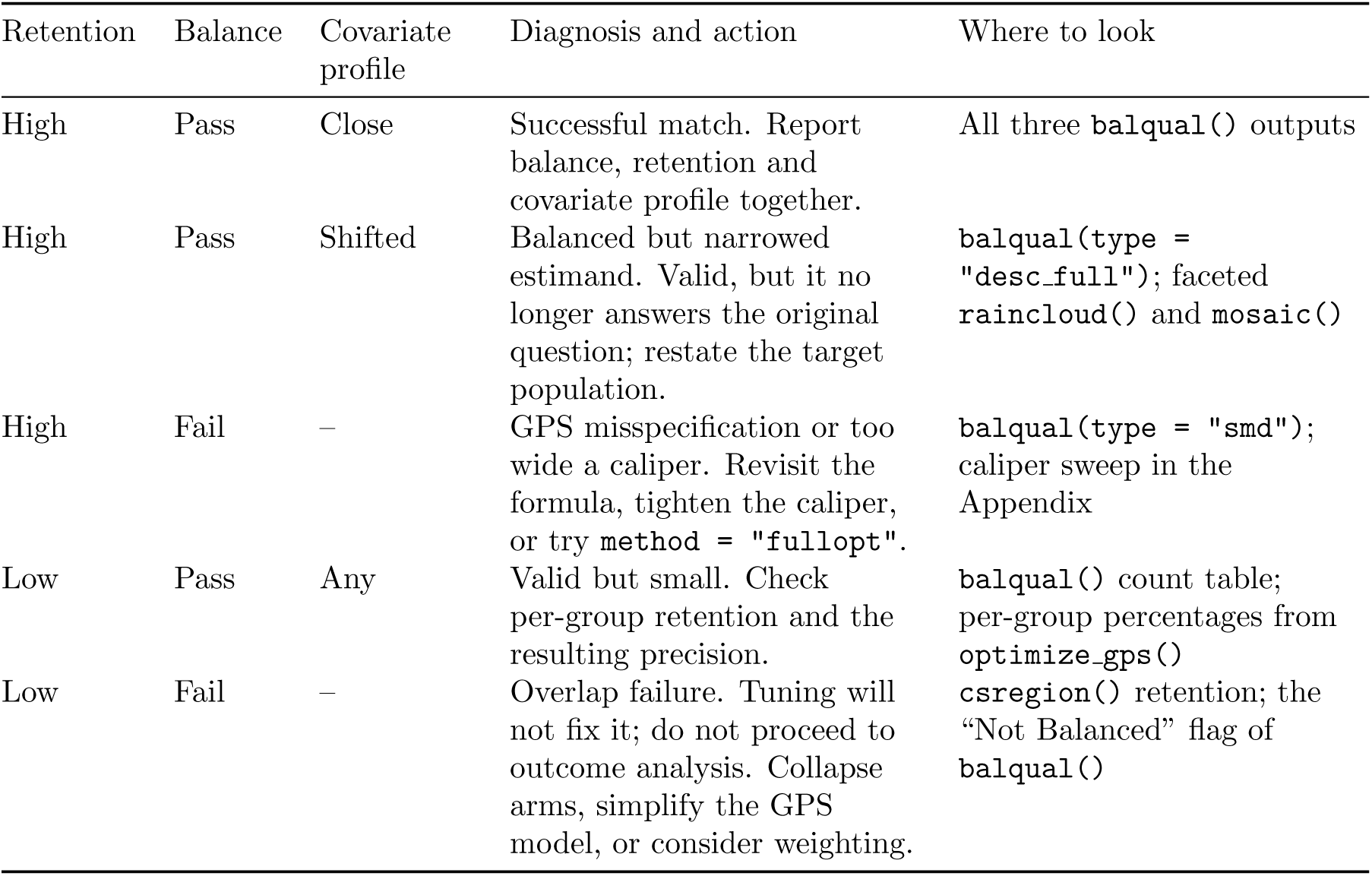
Failure modes of the matching procedure, the combination of diagnostics that identifies each, and the corresponding vecmatch output. A dash indicates that the criterion is not informative once the preceding ones have failed.

| Retention | Balance | Covariate profile | Diagnosis and action | Where to look |
|---|---|---|---|---|
| High | Pass | Close | Successful match. Report balance, retention and covariate profile together. | All three balqual() outputs |
| High | Pass | Shifted | Balanced but narrowed estimand. Valid, but it no longer answers the original question; restate the target population. | balqual(type = "desc_full"); faceted raincloud() and mosaic() |
| High | Fail | – | GPS misspecification or too wide a caliper. Revisit the formula, tighten the caliper, or try method = "fullopt". | balqual(type = "smd"); caliper sweep in the Appendix |
| Low | Pass | Any | Valid but small. Check per-group retention and the resulting precision. | balqual() count table; per-group percentages from optimize_gps() |
| Low | Fail | – | Overlap failure. Tuning will not fix it; do not proceed to outcome analysis. Collapse arms, simplify the GPS model, or consider weighting. | csregion() retention; the “Not Balanced” flag of balqual() |

Two of these rows deserve particular emphasis. The second describes a match that passes both numerical checks while the retained subjects no longer resemble the original sample. This is not a defect of the algorithm, but it does change the question being answered, and it can only be detected by inspecting the covariate descriptions rather than the balance statistics alone.

The last row describes the situation in which the data do not support the intended comparison at all. It is worth understanding why heavy attrition and poor balance tend to occur together. The CSR borders implemented in vecmatch are rectangular, so the retained set is the intersection of the per-group intervals of all *k* treatment groups. The number of observations surviving this step therefore falls as the number of arms grows and as the association between the covariates and treatment assignment strengthens, because a stronger association pushes the estimated GPS values towards zero and one and shrinks the overlap between groups. A dataset in which `csregion()` removes a large majority of the observations and `balqual()` subsequently reports a lack of balance is displaying both symptoms of the same underlying cause, namely that the treatment groups occupy different regions of the covariate space. In this case, the correct conclusion is that the contrast of interest cannot be estimated for the original population without extrapolation, and that at most a contrast for a restricted population remains. Reporting this outcome, rather than tuning the matching parameters until a balanced but uninterpretable subsample emerges, is the scientifically defensible course of action, and the diagnostics described above are intended to make that determination possible.

### 4.3 Conclusions

In summary, vecmatch offers a robust and user-friendly solution for matching across multiple treatment groups. While some loss of observations is inevitable and certain constraints cannot be imposed, the package provides tools to transparently manage these trade-offs. Through its modular architecture and optimization tools, vecmatch enables researchers to conduct valid and reproducible comparisons of multiple groups, which carry a causal interpretation when the identification conditions of Section 1 hold.

## Acknowledgments

We gratefully acknowledge support from the European Union in the form of the ERC Starting Grant EarlyMicroAbs (project number 101075733, awarded to T.V.) and the EU Horizon Health consortium grant ID-DarkMatter-NCD (project number 101136582), coordinated by T.V.

We thank Gabriel Innocenti for his continuous feedback during the development of the package, which greatly contributed to its refinement and improvement.

## Appendix A. Practical considerations: matching quality and percent matched

Researchers using the vecmatch package are often focused on optimizing one of two key attributes of the matching process: matching quality or the percentage of matched samples. However, it is important to note that these two attributes are usually inversely related. Prioritizing higher matching quality typically reduces the number of eligible samples, lowering the percentage of matched samples. Otherwise, maximizing the size of the matched dataset often comes at the expense of matching quality, resulting in a dataset that may not meet desired standards or cutoffs.

To address this issue, it is advisable for researchers to seek a trade-off between these two metrics by adjusting the hyperparameters of the matching process. Below, we provide a quick overview of the key components of the vecmatch package that influence both matching quality and the percentage of matched samples. We also include a brief explanation of how each component impacts these metrics, enabling researchers to understand the underlying logic and enhance their data analysis workflows.

1. **Change the caliper value** - The caliper value defines the range of logit-transformed GPS values for each observation in our dataset, within which it can be matched with other samples. It might be tempting to assume that a wider caliper will automatically lead to a higher number of matched observations but at the cost of lower match quality. This reasoning is valid when using method = “nnm” without replacement. However, based on our experience, for other matching methods and when using matching with replacement, the relationship between caliper width and matching quality may not be strictly linear. Therefore, we encourage researchers to experiment with different caliper values - both small and large - as this parameter can significantly impact the quality of the matching results. A detailed empirical treatment of this relationship, together with practical guidance on selecting a caliper value, is provided in the Appendix section on caliper width (C).
2. **Using different matching methods** - The choice of matching method can significantly impact both matching quality and the percentage of matched observations. Currently, the vecmatch package implements two matching methods for GPS: k- nearest neighbors and full optimal matching. Depending on the specific goal of the matching process - whether prioritizing higher match quality or maximizing the number of retained observations - it is recommended to experiment with both methods in combination with different caliper values to achieve the best results for a given scenario.
3. **Optimizing the formula used to estimate the GPS** - Propensity scores are the core component of the vector matching algorithm, and achieving good matching quality depends on appropriately modeling the relationship between the treatment variable and the confounders. Researchers are thus encouraged to explore these relationships and interactions among various possible confounders to determine the best formulation for estimating the GPS. To identify important confounders, it is advisable to follow the general guidelines for selecting variables in regression models outlined by Gelman (2007). The impact of the correct formula definition on matching quality is generally positive, but its effect on the percentage of matched samples may be ambiguous and needs to be monitored by the researcher. In our experience, more complex formulas often lead to lower matching quality as balancing multiple variables becomes more challenging. Additionally, longer formulas can result in a lower percentage of matched samples, as the added variance from various confounders may reduce the effectiveness of the clustering.
4. **Using different GPS estimation method** - Various methods can be used to estimate the GPS, and this choice can significantly affect matching quality. Different statistical models capture different aspects of the data, sometimes estimating treatment assignment probabilities that differ substantially. Researchers should understand the nature of the data used for GPS estimation and select the appropriate modeling technique. It is also recommended to explicitly specify non-linear relationships in the formula or use a user-defined function for more complex cases. The vecmatch package provides basic multinomial regression models, which should be sufficient in most cases. However, researchers can also explore more advanced methods, such as neural networks or other machine learning techniques, to estimate the GPS. Since it is difficult to predict which model will work best in a given scenario, it is advisable to try multiple approaches and compare the resulting quality.
5. **Optimizing the number of k-means clusters** - While the default number of clusters in Lopez and Gutman (2017) is set to 5, this number can be optimized to improve matching quality. Researchers can determine the optimal number of clusters using methods such as scree plots, the silhouette method, or the gap statistic proposed by Tibshirani et al. (2001).

## Appendix B. Optimizing matching

vecmatch also provides an alternative workflow designed for users focused on maximizing matching quality and sample retention. This workflow leverages a random search algorithm to explore the parameter space and identify optimal combinations based on user-defined objectives. The process consists of four steps:

1. **Define the search space** - The matching process involves numerous parameters, but users typically wish to optimize only a subset relevant to their specific goals. The `make_opt_args()` function is used to define and validate this subset of parameters or parameter vectors. It prepares the search space for the main optimization routine and prints a summary to the console, including the number of possible parameter combinations:

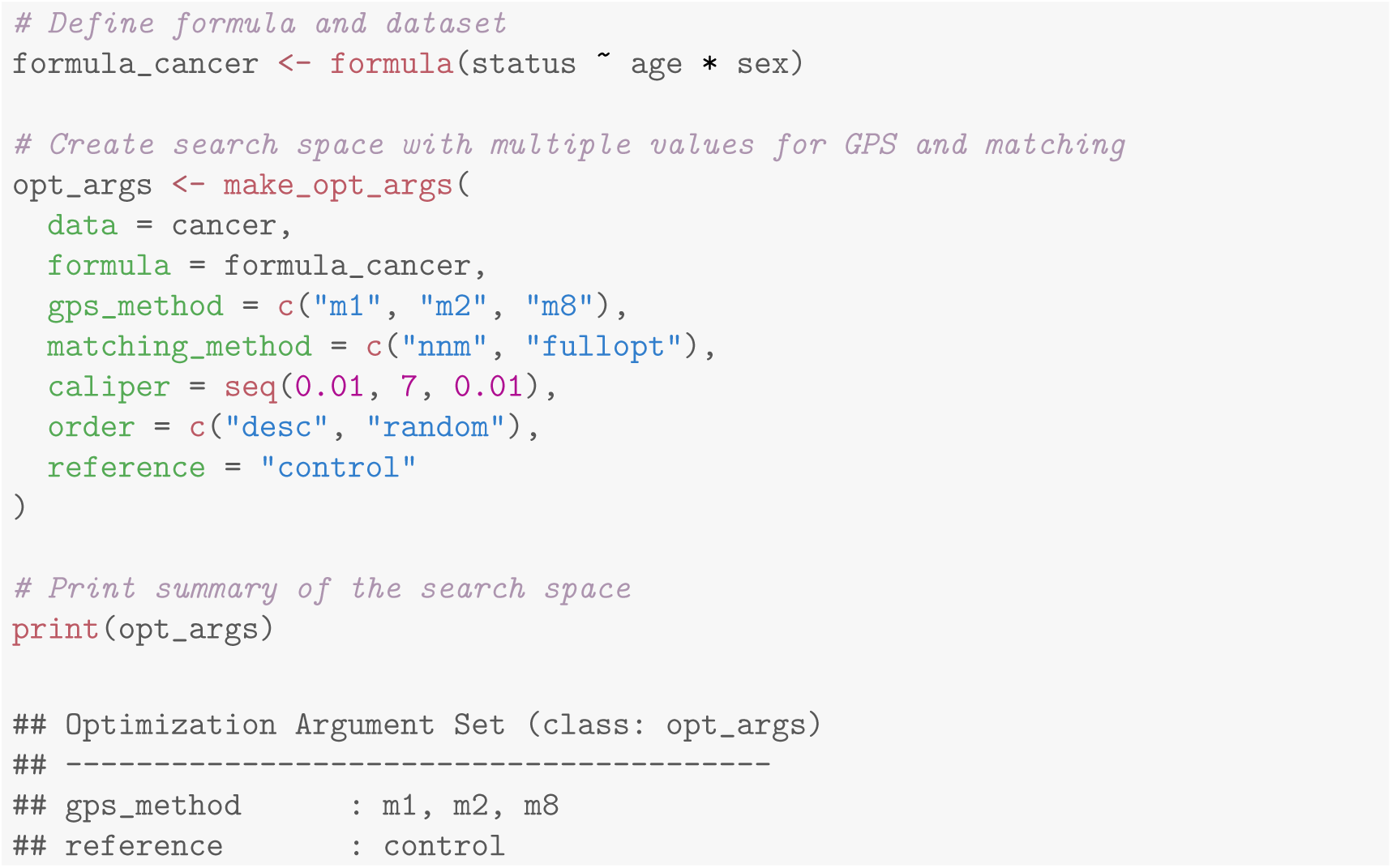
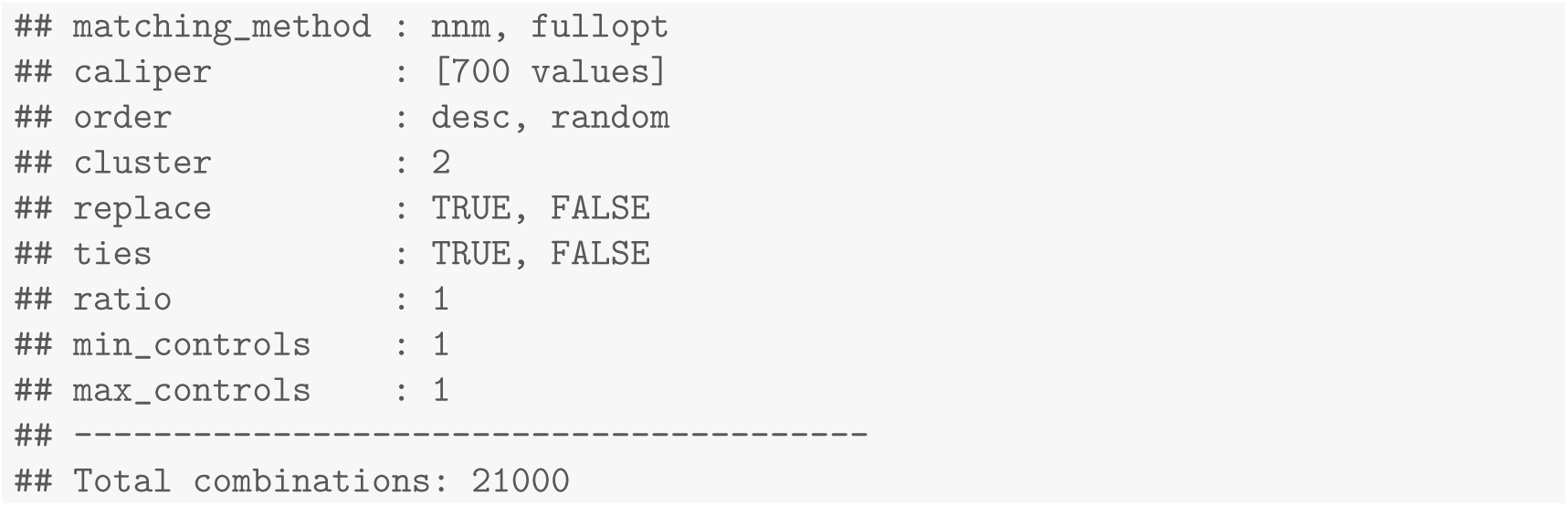 The `make_opt_args()` function includes a set of default parameters, which are documented in the official package manual. In the example case, the defined parameter space results in a total of 21,000 possible combinations. Once the search space has been constructed, it is passed to the main optimization function in the next step of the workflow.
2. **Optimization function** - This function serves as the core component of the optimization workflow. It performs Steps 2-4 of the main `vecmatch` procedure (excluding the initial assessment of covariate imbalance, as it is not necessary here) for a user- defined number of parameter combinations. For each run, it calculates the overall maximal standardized mean difference (SMD), which serves as the primary matching quality metric in `vecmatch`. Additionally, it computes partial SMDs for all pairwise treatment group comparisons and for all variables specified in the `formula` argument - including both main effects and interactions. The function also estimates the percentage of matched observations per treatment group. These percentages are derived from `balqual()` and are calculated relative to the number of observations remaining after CSR filtering, rather than relative to the original sample size. This allows users to filter and prioritize results based on group-specific matching retention. To improve performance, the optimization function supports multithreading. Because large-scale parameter searches can be computationally intensive, especially with many iterations, parallel processing can significantly reduce runtime. However, users should be aware that multithreading in vecmatch can impose high memory demands, particularly with large datasets. Each parallel R session consumes memory independently, which may lead to memory overload and system crashes if not properly managed. It is therefore essential to carefully control the number of threads and monitor available system memory during execution.

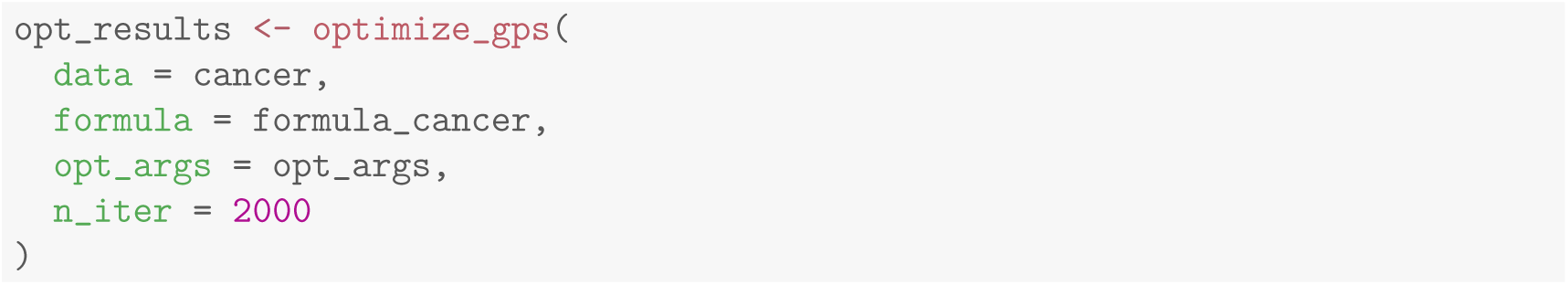
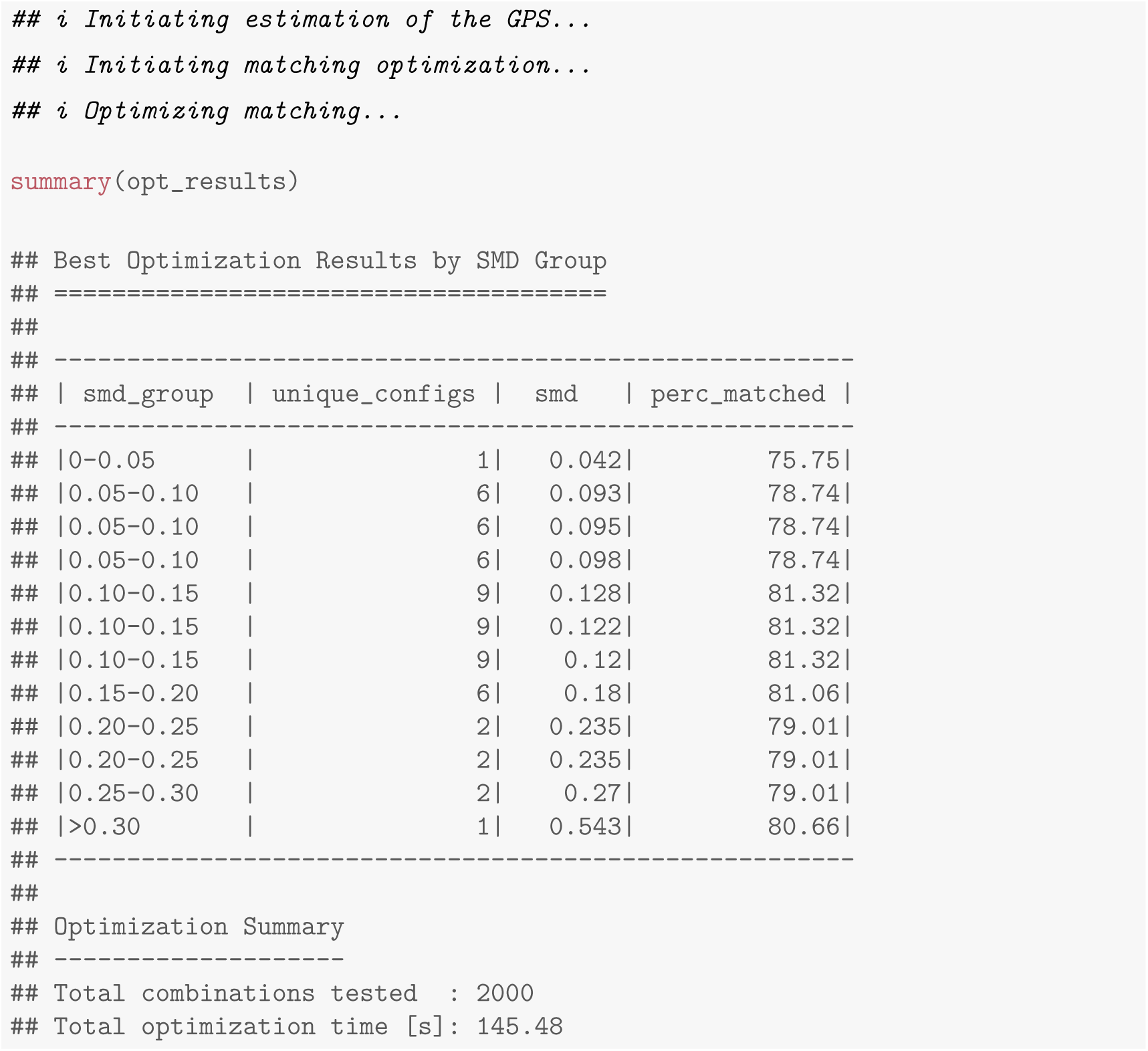 The output of the optimization function can be passed to the `select_opt()` function, which assists in identifying the optimal combination of matching parameters based on the researcher’s predefined objectives.
3. **Selecting the Results** - Researchers may have varying objectives in the matching process. For instance, they might aim to retain the maximum number of observations in specific treatment groups (e.g., a control group), while placing less emphasis on others. The `select_opt()` function enables filtering and selection of the optimization results based on such user-defined priorities. For example, to prioritize maximizing the number of matched observations in the control group, one would use the following code:

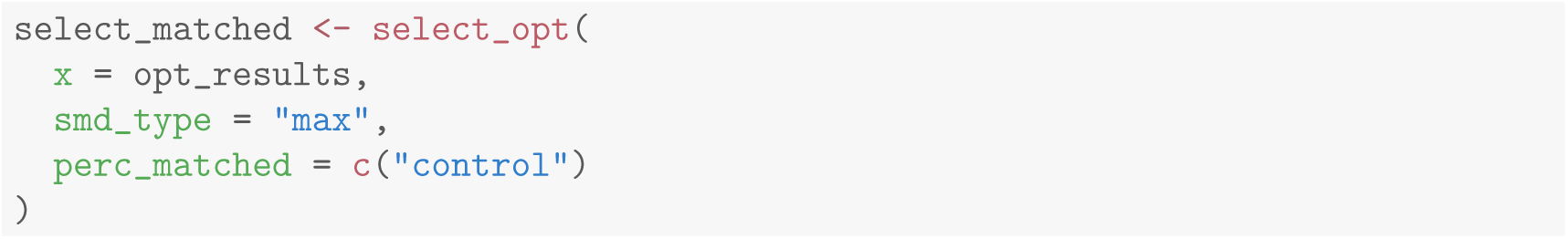
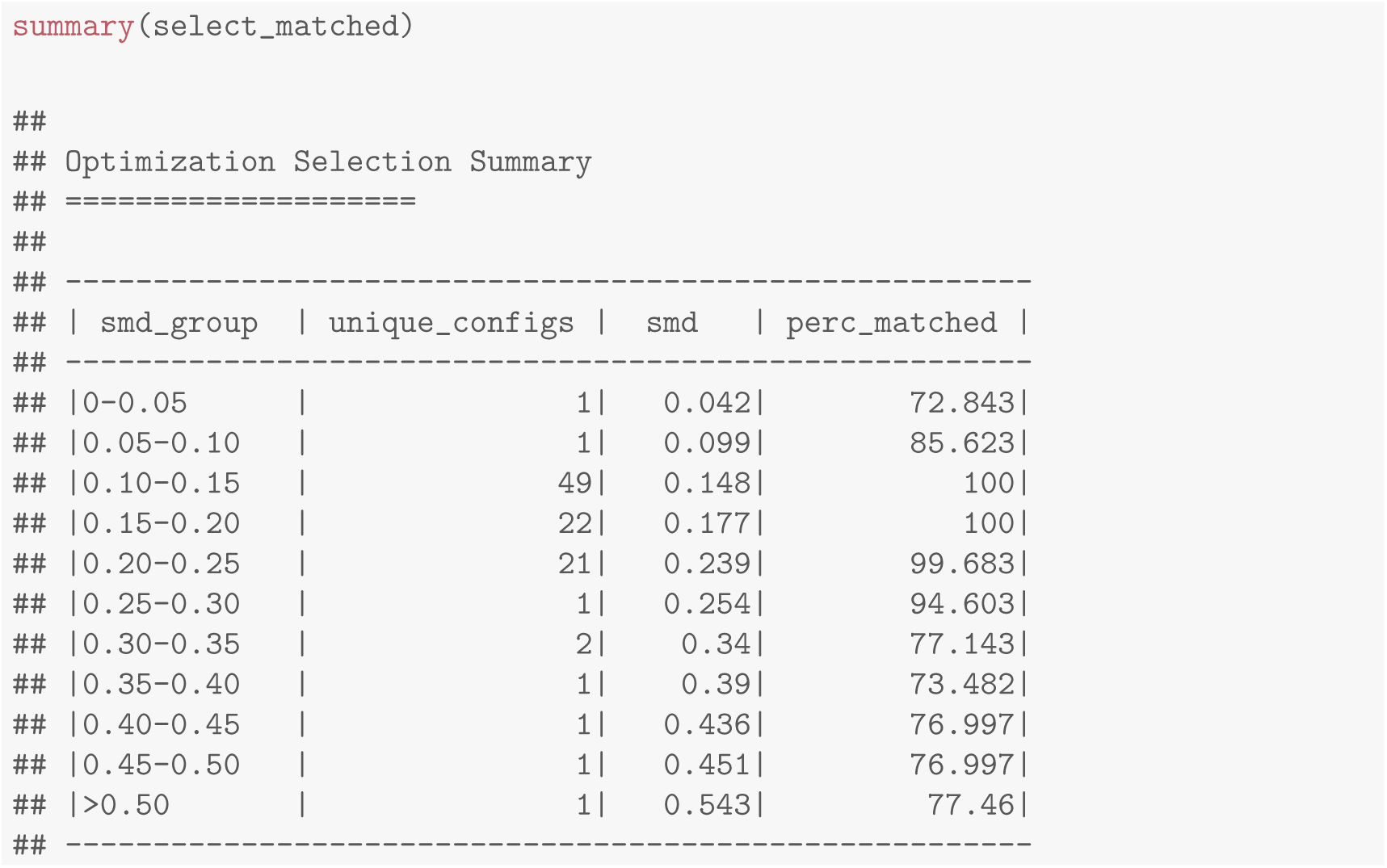 The table lists, for each band of the maximum SMD, the best retention achieved within it. Applying the selection rule of the Appendix section on choosing a caliper (C) - fix the balance criterion first, then maximize retention among the configurations that satisfy it - the relevant row is the one below the strict cutoff of 0.10, which reaches a maximum SMD of 0.099 while matching roughly 86 percent of the control group. Configurations retaining all control observations exist, but only at a maximum SMD of about 0.15, above the threshold we recommend; reporting the whole table rather than a single winner is what makes that trade-off visible to the reader.
4. **Refitting** - After completing the optimization workflow, users may wish to obtain the final matched dataset. To do this, a manual refitting step is required. The selected set of optimal parameters must be extracted from the output of the optimization or selection functions, and Steps 2-4 of the main vecmatch workflow must be executed manually using these parameters.

## Appendix C. Caliper width and matching quality

Among the parameters of `match_gps(), the caliper` is the one most directly under the researcher’s control and the one with the most immediate effect on both matching quality and sample retention. In vecmatch, the caliper is not expressed on the raw GPS scale: it defines the allowable matching distance as a fraction of the standard deviation of the logit-transformed GPS. The package default, `caliper` = 0.2, therefore corresponds to the conventional rule of thumb of 0.2 standard deviations of the logit propensity score, originally proposed by Rosenbaum and Rubin (1985) and later supported by the simulation work of Austin (2011), who found caliper widths in the range of 0.2 standard deviations to perform well across a variety of binary-treatment scenarios.

This convention is a reasonable starting point, but it should not be treated as a value to be adopted without checking. The published guidance was derived for the binary-treatment setting, whereas vector matching performs a sequence of pairwise matches within *k*-means clusters, so the effective constraint imposed by a given caliper depends on the number of clusters, the matching method, and the shape of the GPS distribution. At first glance, it might seem that the relationship between matching quality and caliper width is linear, or at least monotone. Narrowing the caliper restricts the range of logit-transformed GPS values within which an observation may be matched, which should in theory produce two effects: an improvement in matching quality, since only observations that are close to one another are paired, and a decrease in the number of matched observations, since fewer candidates fall within the tighter range. As noted in the Appendix section on practical considerations (A), however, such a relationship is clearly visible only in simple matching scenarios using `method = “nnm”` without replacement, and only for low caliper values. When other matching methods are applied, the nature of this relationship can change substantially and deviate from a monotone pattern.

To make this concrete, we examined the relationship between the caliper, matching quality, and the percentage of matched observations across several matching scenarios using the `cancer` dataset shipped with `vecmatch`. This dataset is well suited to the purpose because `age` is a genuinely continuous, approximately normally distributed variable. Datasets in which the continuous confounder is recorded on a coarse grid - for instance, age rounded to whole years in a registry - produce many observations with identical covariate patterns and hence identical GPS values, which masks the effect of the caliper: below a certain width the set of eligible partners stops changing, and the sweep becomes uninformative. Researchers running the analysis described here on their own data should keep this limitation in mind.

GPS values were estimated using the “multinom” method with sex, age, and their interaction as confounders, and the control group as the reference category. Five specifications of match `gps()` were then compared:

1. **Model 1:** full optimal matching (`method = “fullopt”`),
2. **Model 2:** *k*-nearest neighbors matching (KNNM) without replacement (`method = “nnm”, replace = FALSE`),
3. **Model 3:** KNNM without replacement with ascending sorting order (`method = “nnm”, replace = FALSE, order = “asc”`),
4. **Model 4:** KNNM with replacement (`method = “nnm”, replace = TRUE`),
5. **Model 5:** KNNM with replacement and ties retained (`method = “nnm”, replace = TRUE, ties = TRUE`).

All remaining parameters were left at their default values, and the number of *k*-means clusters was fixed at 2 across all scenarios. For each specification, 150 caliper values were tested, ranging from 0.01 to 1.50 in increments of 0.01.

The results are presented in Figures 6 to 8. Figure 6 relates the caliper value to matching quality, represented by the highest SMD in the matched dataset. Figure 7 shows the corresponding relationship with the percentage of matched observations. Finally, Figure 8 plots the maximum SMD values against their corresponding percentages of matched observations, with regression lines fitted using generalized additive models.

**Figure 6:**
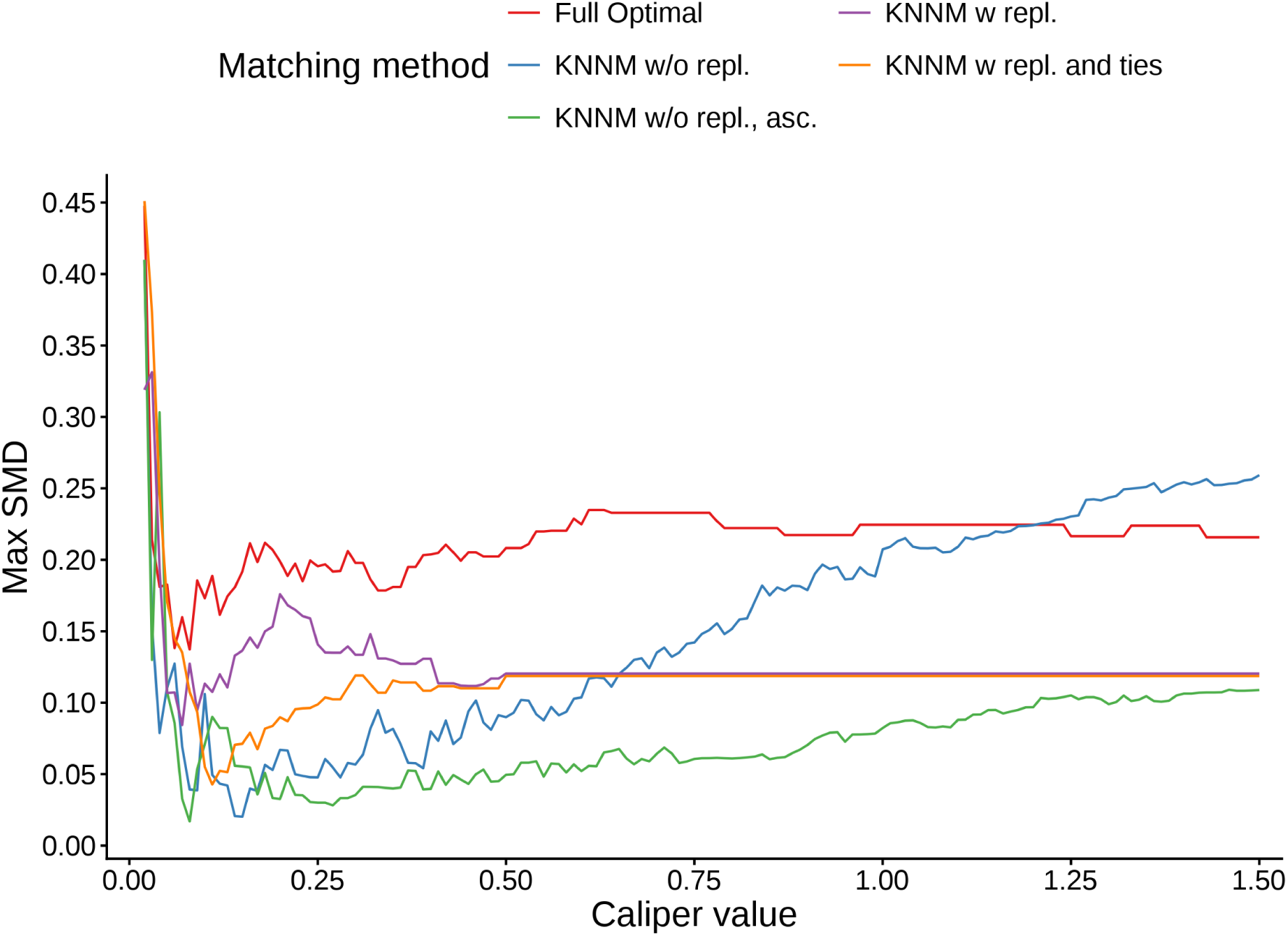
The relationship between the value of the caliper parameter and the quality of matching, expressed as the maximum standardized mean difference (SMD) in the matched dataset, for five matching specifications. Lower values on the vertical axis indicate better balance. A monotone increase in the maximum SMD with increasing caliper width is visible only for the two k-nearest neighbors specifications without replacement.

**Figure 7:**
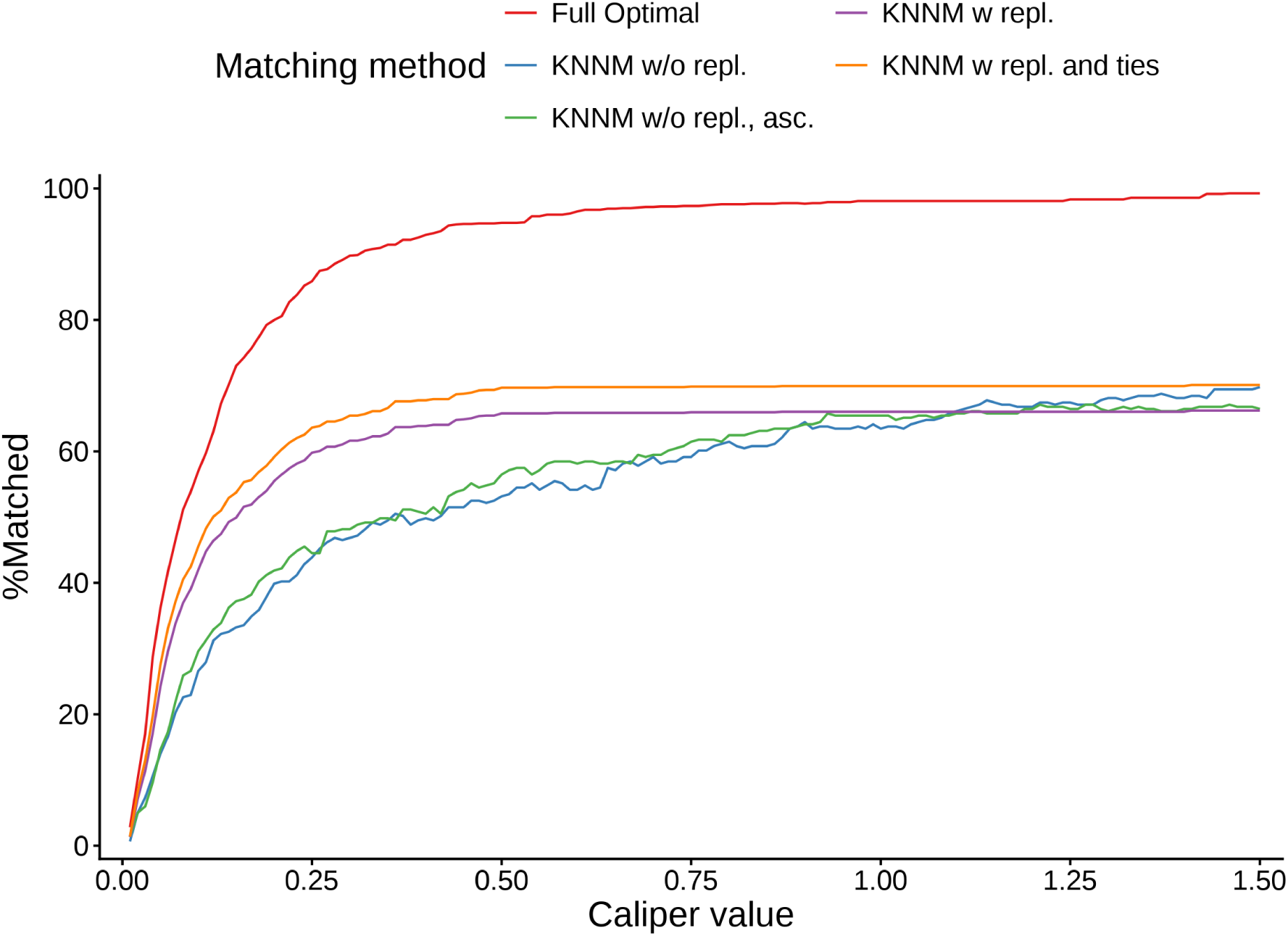
The relationship between the value of the caliper parameter and the percentage of matched observations for five matching specifications. In contrast to the matching quality shown in the preceding figure, the trend here is consistent across all methods and resembles a logarithmic curve that saturates once the caliper stops being the binding constraint.

In Figure 6, a clear trend is observed only for KNNM without replacement. For these two specifications the maximum SMD grows with the caliper, gently up to roughly caliper = 0.5 and steeply thereafter for the default descending order, and sorting the GPS values in ascending order generally yields better results - that is, lower SMD values - than the default descending order. For the remaining methods, the relationship between caliper and matching quality is considerably harder to characterize. For KNNM with replacement, the trend depends in part on the value of the ties argument: the two curves differ markedly at small caliper widths and converge only from roughly caliper = 0.5 onward, where both settle at a maximum SMD of about 0.12. Full optimal matching produces the highest SMD values over most of the range and never falls below 0.137, so the conventional threshold of 0.10 is out of reach for this method on these data. Its curve is also steep and irregular in the region around the package default, so a researcher who accepted the default without inspecting its neighborhood would have no way of knowing whether they had landed on a favorable value or a fragile one.

The overall trend is much easier to identify for the percentage of matched observations in Figure 7. Increasing the caliper had a consistently positive effect on the number of matched observations for every method, following a curve resembling a logarithmic function. Full optimal matching achieved the highest retention throughout, matching approximately 90 percent of the dataset at a caliper of about 0.30 and exceeding 97 percent for caliper values above 0.65, beyond which the curve is essentially flat. KNNM with replacement performed less well, plateauing at roughly 66 percent, with ties = TRUE yielding a slight increase over the default. The weakest retention was observed for both variants of KNNM without replacement, at least for caliper values below approximately 1.10; for larger calipers their performance became comparable to KNNM with replacement.

Figure 8 provides the most direct way of reading the trade-off, since it removes the caliper from the axes entirely and shows only the balance and retention that a given caliper buys. It makes clear that the best method depends on the balance one insists on. If the goal is to achieve post-matching SMD values below the conventional cutoff of 0.10, full optimal matching is not an option at all on these data, and the strongest choice is KNNM without replacement with the GPS sorted in ascending order, which retains about 67 percent of the dataset at that level of balance. Under a more liberal threshold of 0.20, full optimal matching becomes the best-performing option, with retention close to 95 percent. The plot also makes the cost of the wrong combination visible: the KNNM specifications with replacement scatter widely, with many caliper values yielding both poor balance and low retention simultaneously.

**Figure 8:**
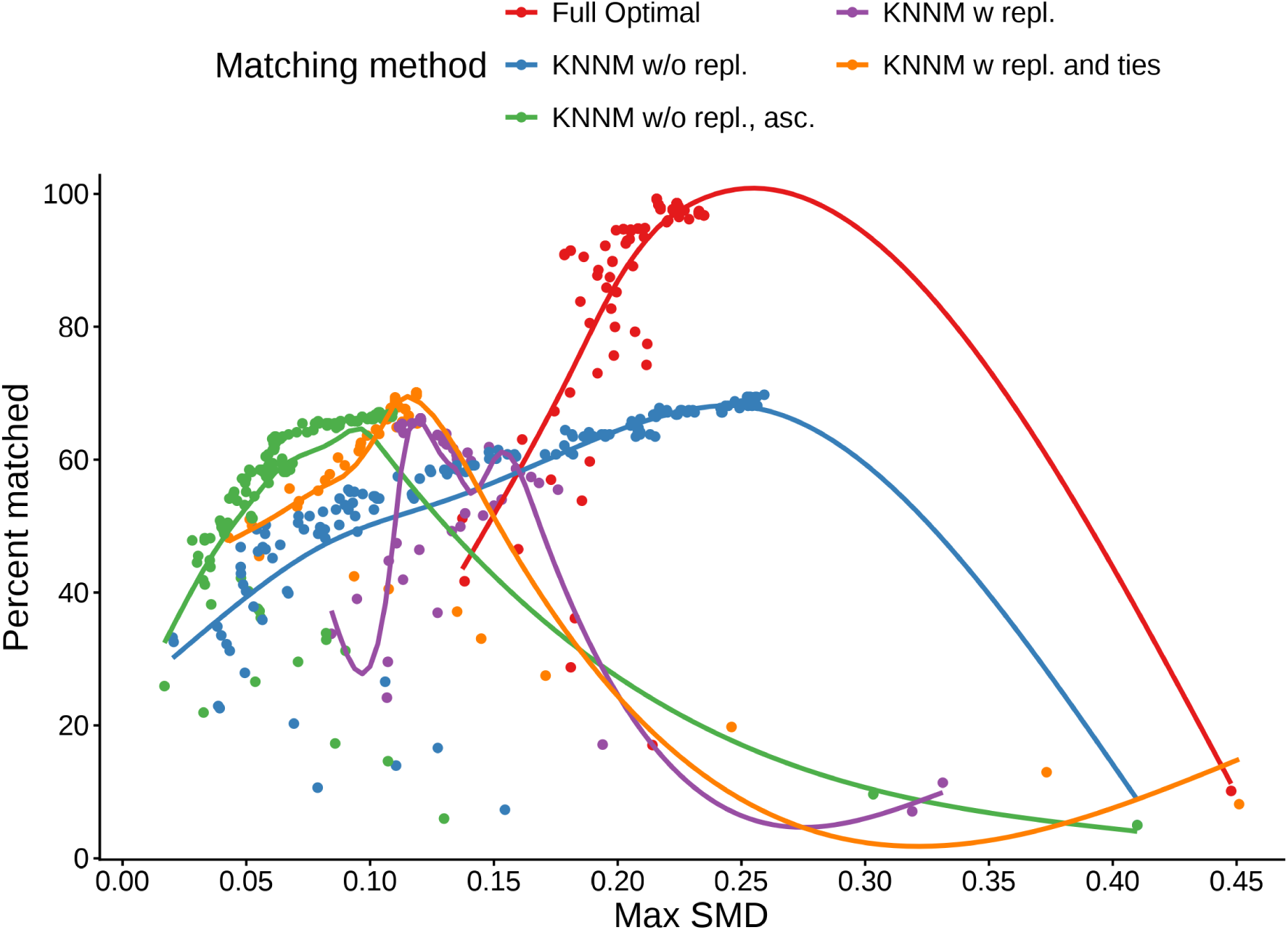
The relationship between maximum standardized mean difference (SMD) values and the corresponding percentages of matched observations for different matching methods, visualized using color-coded points. Each point corresponds to one caliper value from the sweep. For each matching method, the best-fitting curve was estimated using generalized additive models and plotted on the graph. These regression curves serve only as a visual aid to illustrate the overall trend in the data and should be interpreted with caution due to the high dispersion of observations in some of the examined groups.

### C.1 Choosing a caliper value

The analysis above is illustrative rather than prescriptive. The results depend on the dataset, the formula used for GPS estimation, the matching method, and the number of *k*-means clusters, and they should not be transferred directly to another analysis. What does generalize is the procedure. Based on the patterns shown above, we suggest the following approach to selecting a caliper.

1. **Start from the default.** Fit an initial model with caliper = 0.2, the package default and the conventional 0.2 standard deviation rule (Rosenbaum and Rubin, 1985; Austin, 2011), and record the maximum SMD and the percentage of matched observations reported by `balqual()`. This establishes a reference point but should not be the final answer.
2. **Sweep a coarse grid before committing.** Repeat the matching over a range of caliper values - a grid from 0.01 to roughly 1.5 in steps of 0.01 is sufficient in most applications, and coarser steps are adequate for a first pass - holding all other parameters fixed, and record both metrics at each value. Because the relationship is not monotone for most methods, a sweep is the only reliable way to find out how the caliper behaves on a given dataset. This is also the reason we recommend against reporting a single caliper value without any indication of its stability.
3. **Read the trade-off, not the caliper.** Plot the maximum SMD against the percentage of matched observations, as in Figure 8, rather than plotting either quantity against the caliper alone. The caliper is a means to an end, and two very different caliper values can produce nearly identical balance and retention.
4. **Fix the balance cutoff, then maximize retention.** Decide the acceptable maximum SMD in advance - 0.10 is the conventional strict choice and 0.20 the more liberal one - and among the caliper values that satisfy it, choose the one maximizing retention. Prefer values situated in a flat, stable region of the curve over isolated minima, since the latter are likely to reflect the particular random seed rather than a genuine property of the data.
5. **Repeat across matching methods.** As Figures 6 and 7 show, the best caliper for one method is not the best for another, and the ranking of methods itself changes with the caliper. The caliper should therefore be tuned jointly with the matching method rather than in isolation.

Researchers who prefer not to implement this sweep manually can use the optimization workflow described in the Appendix section on optimizing matching (B). Passing a caliper vector to `make_opt_args()` and running `optimize_gps()` performs the same search automatically, jointly with the other matching parameters, and `select_opt()` then applies the selection rule described in step 4 above to the accumulated results.

